# Cryptic Genes for Interbacterial Antagonism distinguish *Rickettsia* species Infecting Blacklegged Tick from other *Rickettsia* pathogens

**DOI:** 10.1101/2022.02.21.481368

**Authors:** Victoria I. Verhoeve, Tyesha D. Fauntleroy, Riley G. Risteen, Timothy P. Driscoll, Joseph J. Gillespie

## Abstract

**Background:** Genus *Rickettsia* (*Alphaproteobacteria*: Rickettsiales) encompasses numerous obligate intracellular species with predominantly ciliate and arthropod hosts. Notable species are pathogens transmitted to mammals by blood-feeding arthropods. Mammalian pathogenicity evolved from basal, non-pathogenic host-associations; however, some non-pathogens are closely related to pathogens. One such species, *Rickettsia buchneri*, is prevalent in the blacklegged tick, *Ixodes scapularis*. While *I*. *scapularis* transmits several pathogens to humans, it does not transmit *Rickettsia* pathogens. We hypothesize that *R*. *buchneri* established a mutualism with *I*. *scapularis*, blocking tick superinfection with *Rickettsia* pathogens.

**Methods:** To improve estimates for assessing *R*. *buchneri* infection frequency in blacklegged tick populations, we used comparative genomics to identify an *R*. *buchneri* gene (*REIS_1424*) not present in other *Rickettsia* species present throughout the *I*. *scapularis* geographic range. Bioinformatic and phylogenomics approaches were employed to propose a function for the hypothetical protein (263 aa) encoded by *REIS_1424*.

**Results:** REIS_1424 has few analogs in other Rickettsiales genomes and greatest similarity to non-Proteobacteria proteins. This cohort of proteins varies greatly in size and domain composition, possessing characteristics of Recombination hotspot (Rhs) and contact dependent growth inhibition (CDI) toxins, with similarity limited to proximal C-termini (∼145 aa). This domain was named CDI-like/Rhs-like CT toxin (CRCT). As such proteins are often found as toxin-antidote (TA) modules, we interrogated REIS_1423 (151 aa) as a putative antidote. Indeed, REIS_1423 is similar to proteins encoded upstream of CRCT domain-containing proteins. Accordingly, we named these proteins CDI-like/Rhs-like C-terminal toxin antidotes (CRCA). *R*. *buchneri* expressed both *REIS_1423* and *REIS_1424* in tick cell culture, and PCR assays showed specificity for *R*. *buchneri* over other rickettsiae and utility for positive detection in three tick populations. Finally, phylogenomics analyses uncovered divergent CRCT/CRCA modules in varying states of conservation; however, only *R*. *buchneri* and related Tamurae/Ixodes Group rickettsiae carry complete TA modules.

**Conclusion:** We hypothesize that *Rickettsia* CRCT/CRCA modules circulate in the *Rickettsia* mobile gene pool, arming rickettsiae for battle over arthropod colonization. While its functional significance remains to be tested, *R*. *buchneri* CRCT/CRCA serves as a marker to positively identify infection and begin deciphering the role this endosymbiont plays in the biology of the blacklegged tick.

## Introduction

The blacklegged tick (*Ixodes scapularis*), more commonly referred to as deer tick, is of vital importance to human health as a vector of several infectious disease agents: e.g., *Borrelia* species (Lyme disease), *Anaplasma phagocytophilum* (anaplasmosis), *Babesia* and *Theileria* parasites (babesiosis, theileriosis), and Powassan Flavivirus (Powassan disease) (Wormser et al., 2006; Madison-Antenucci et al., 2020). Curiously, blacklegged tick does not infect humans with *Rickettsia* pathogens, despite overlapping in geographic range with other tick species that do; e.g., American dog tick (*Dermacentor variabilis*), Brown dog tick (*Rhipicephalus* sanguineus), Gulf Coast tick (*Amblyomma maculatum*), and Lone Star tick (*Amblyomma americanum*) (Walker and Ismail, 2008; Lee et al., 2014; Sanchez-Vicente et al., 2019). However, *I*. *scapularis* is predominantly infected with a species of *Rickettsia*, *R*. *buchneri*, that is considered a non-pathogen of humans and has not been detected in vertebrates (Kurtti et al., 2015). The presence of *R*. *buchneri* in tick ovaries (Munderloh et al., 2005), high infection rate in ticks regardless of co-infection with other intracellular bacteria (Billings et al.; Magnarelli et al., 1991; Benson et al., 2004; Swanson and Norris, 2007; Troughton and Levin, 2007; Steiner et al., 2008) or composition of other microbiota (Moreno et al., 2006; Narasimhan et al., 2014; van Treuren et al., 2015; Abraham et al., 2017; Ross et al., 2018; Thapa et al., 2019; Tokarz et al., 2019) hint at an underappreciated host-microbe relationship in need of further investigation.

A decade ago, we reported the first genome sequence of *R*. *buchneri* by assembling bacterial-like sequencing reads generated by the *I*. *scapularis* genome project (Gillespie et al., 2012a; Gulia-Nuss et al., 2016). Prior to this, phylogenomics analyses of diverse *Rickettsia* genomes indicated high conservation in synteny, moderate pseudogenization, one or zero plasmids, and relatively few **mobile genetic elements** (**MGEs**) (Darby et al., 2007; Fuxelius et al., 2008; Gillespie et al., 2008, 2012b). Several attributes emerged from our analyses highlighting the oddity of the *R. buchneri* genome, including 1) pronounced pseudogenization relative to other rickettsiae, especially for genes in other **Spotted Fever Group** (**SFG**) rickettsiae with characterized functions in vertebrate pathogenesis, 2) a substantial number of transposases (∼30% of total CDS), 3) four novel plasmids (pREIS1-4), and 4) nine copies (seven chromosomal and two plasmid) of the **Rickettsiales Amplified Genetic Element** (**RAGE**), a conjugative transposon found as single-copy in certain other *Rickettsia* genomes (Ogata et al., 2006; Blanc et al., 2007). Estimated phylogenies placed *R*. *buchneri* basal to all SFG rickettsiae, indicating a substantially different evolutionary track relative to derived SFG rickettsiae lineages, as well as species in the **Typhus Group** (**TG**) and **Transitional Group** (**TRG**) rickettsiae, yet inordinate **lateral gene transfer** (**LGT**) with the ancestral *R*. *bellii* and other intracellular bacteria. This was exemplified by *R*. *buchneri’s* RAGEs, which encode numerous genes with functions critical for obligate intracellular life, leading to our hypothesis that RAGEs and other MGEs are vehicles for gene acquisitions that offset high rates of pseudogenization (Gillespie et al., 2012a) (**Fig. 1A**).

**FIGURE 1.**
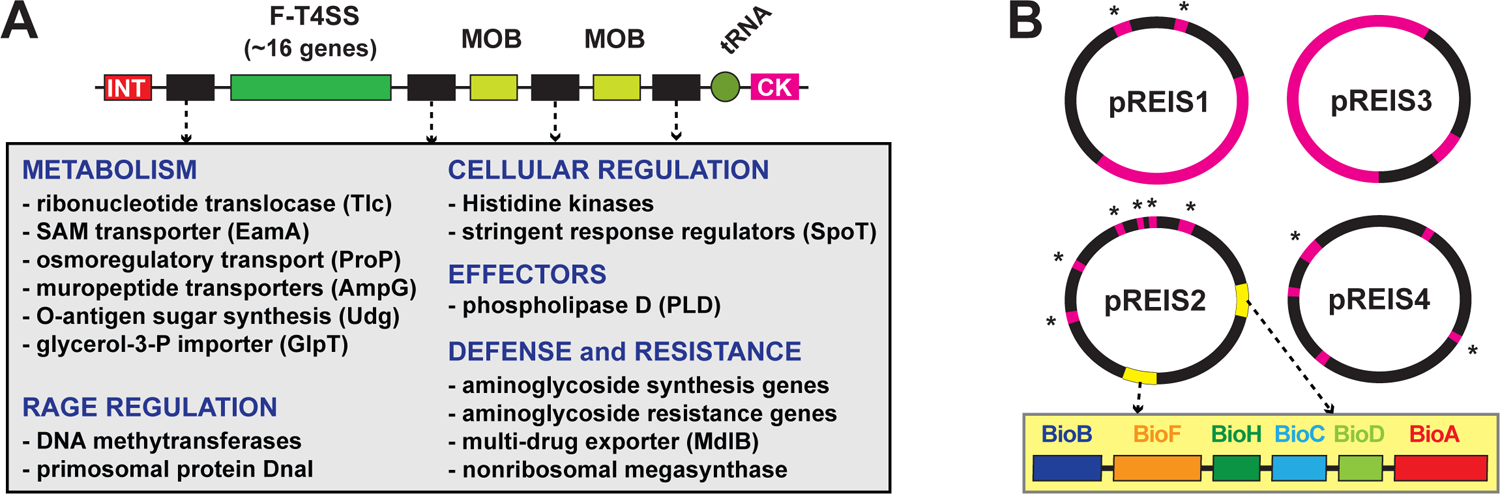
Two extraordinary features of the *R*. *buchneri* genome include RAGEs and four unique plasmids (pREIS1-4) (Gillespie et al., 2012a). (**A)** General schema of RAGEs: INT, integrase; MOB, mobilization genes (*traT* and *traAI*). RAGEs typically insert in tRNA genes near cytosine kinase (CK)-encoding loci. Cargo genes (black) occur in variable clusters. Grey inset lists recurring cargo genes grouped by class. (**B**) Schema of pREIS plasmids: pink, full-length RAGE; pink w/ asterisk, MOB genes; yellow, BOOM (illustrated in yellow inset). BOOM genes are uniquely arrayed relative to other bacteria that carry biotin synthesis genes (Gillespie et al., 2012a)

The *R*. *buchneri* genome was also found to encode many genes absent from other rickettsiae. Noteworthy are those encoding **aminoglycoside antibiotic** (**AGAB**) synthesis/resistance genes and **polyketide synthase** (**PKS**)-containing **nonribosomal protein synthases** (**NRPS**), hinting at a defense arsenal of antibiotics and 2° metabolites. Furthermore, plasmid pREIS2 of *R*. *buchneri* carries identical duplications of the **biotin synthesis operon of obligate microbes** (**BOOM**), (**Fig. 1B**), which is only found in a limited range of obligate intracellular species, including certain wolbachiae (Nikoh et al., 2014; Gerth and Bleidorn, 2017; Balvín et al., 2018; Driscoll et al., 2020), *Cardinium* (Penz et al., 2012; Zeng et al., 2018) and *Legionella* (Ríhová et al., 2017) species. As some wolbachiae provide biotin to their insect hosts (Nikoh et al., 2014; Ju et al., 2019), *R*. *buchneri* may supplement biotin to the blacklegged tick bloodmeal, which is a poor source of B vitamins (Lehane and Lehane, 2010; Manzano-Marín et al., 2015). This possibility, coupled with potential to provision blacklegged tick with antibiotics and 2° metabolites, indicates *R*. *buchneri* has characteristics of a mutualistic bacterium unlike any rickettsiae heretofore analyzed from a genomic perspective.

If *R*. *buchneri* is a mutualist of blacklegged tick, infection frequency should be very high in nature. Reported *R*. *buchneri* infection rates in *I*. *scapularis* populations range from under 20% to over 80%; however, this disparity may reflect different sampling strategies not aiming to directly detect *R*. *buchneri* or distinguish this species from other rickettsiae (i.e., using general *Rickettsia* PCR primers). Differences in tissue sampling across studies could also distort accuracy in detecting *R*. *buchneri* infection if ovaries are not sampled. Furthermore, infection rates are higher in females and nymphs (Cross et al., 2018; Hagen et al., 2018; Gil et al., 2020), indicating a reduction in males after molting to adulthood; thus, sexing is important to understand true infection frequency.

In this report, we developed a *R*. *buchneri*-specific PCR assay to help rule out *Rickettsia* co-infections in future studies. Our strategy was guided by re-evaluating a set of genes previously determined to be unique to *R*. *buchneri* (Gillespie et al., 2012a). One gene, *REIS-1424* (encoding a 263 aa hypothetical protein), was shown by *in silico* analysis to be absent in other rickettsiae known throughout the geographic range of *I*. *scapularis*. Unexpectedly, bioinformatic and phylogenomics analyses indicated that *REIS_1424* and its neighboring gene *REIS_1423* comprise a **toxin-antidote** (**TA**) module typical of certain bacterial systems used primarily for interbacterial warfare. As with numerous other *R*. *buchneri*-specific genes, *REIS_1424* and *REIS_1423* show evidence of LGT from distantly related non-proteobacteria. Similar cryptic TA modules are recurrent in rickettsiae and illuminate a potential mechanism for *Rickettsia* competition in arthropod hosts.

## Materials and Methods

### Identifying a *R*. *buchneri*-specific gene

A set of genes (739 singletons) from our prior report lacked significant similarity to genes in other *Rickettsia* genomes (Gillespie et al., 2012a); accordingly, these were used as queries in Blastn searches against the NCBI ‘Rickettsia’ databases (taxid 780). Genes with significant similarity to genes from rickettsiae present within the geographic distribution of *I*. *scapularis* were removed. Further, we excluded genes encoded on plasmids, those present within chromosomal RAGES, and those encoding transposases or related elements. The remaining genes were then evaluated using bioinformatics analysis to determine the likelihood that they encode functional proteins.

### Compiling toxin and antidote datasets

For toxins, proximal (∼100 aa) C-terminal sequences of *R*. *buchneri* protein REIS_1424 (EER22217) and another rickettsial species “*Candidatus* Jidaibacter acanthamoeba” str. UWC36 protein NF27_IC00050 (KIE04387) were used as queries in Blastp searches to compile and analyze diverse proteins harboring significant similarity across complementary proximal C-terminal sequences. For antidotes, entire sequences for *R*. *buchneri* protein REIS_1423 (EER22217 with an adjusted start site adding 41 aa at the N-terminus) and “*Candidatus* Jidaibacter acanthamoeba” str. UWC36 protein NF27_IC00040 (KIE04386) were used as queries in Blastp searches to compile and analyze diverse proteins harboring significant similarity across the entire lengths of the queries. Analyses utilized our HaloBlast method, which is a combinatorial Blastp-based approach for interrogating proteins for LGT (Driscoll et al., 2013). Individual Blastp searches were conducted against five distinct taxonomic databases: 1) “Rickettsia” (NCBI taxid 780)”, 2) “Rickettsiales” (taxid: 766) excluding “Rickettsia”, 3) “Alphaproteobacteria” (taxid: 28211) excluding “Rickettsiales”, 4) “Proteobacteria” (taxid: 1224) excluding “Alphaproteobacteria”, 5) “Bacteria” (taxid: 2) excluding Proteobacteria”, and 6) “minus bacteria”). All subjects from each search were ranked by *Sm* score (= *b* * *I* * *Q*, where *b* is the bitscore of the match, *I* is the percent identity, and *Q* is the percent length of the query that aligned), a comparative sequence similarity score designed to de-emphasize highly significant matches to short stretches of the query in favor of longer stretches of similarity (Driscoll et al., 2013). The “halos” or separate database searches were then compared to one another to determine the taxon with the strongest similarity to the query sequences.

### Toxin characterization

HaloBlast subjects from the searches with REIS_1424 and NF27_IC00050 as queries were analyzed in two ways. First, only sequences matching the proximal (∼100 aa) C-terminal sequences of the query were compiled and aligned with MUSCLE using default parameters (Edgar, 2004). The entire alignment was then visualized as sequence logos using WebLogo (Crooks et al., 2004). Second, two representative sequences per halo were selected for domain predictions across the entire protein. EMBL’s **Simple Modular Architecture Research Tool** (**SMART**) (Letunic and Bork, 2017) and /or the **Protein Homology/analogY Recognition Engine** V 2.0 (**Phyre2**) (Kelley and Sternberg, 2009) were used to predict and evaluate the following domains: UBA (ubiquitin-associated) (Mueller and Feigon, 2002); haemagglutination activity site (Kajava et al., 2001); hemagglutinin repeats (Pfam ID PF13332); Peptidase M43 domain (Rawlings and Barrett, 1995); endonuclease III (Bruner et al., 2000); RHS repeat (Busby et al., 2013); VENN motif (Aoki et al., 2010; Zhang et al., 2011); DUF637: hemagglutinin-/hemolysin-associated domain (PF04830); alanine-rich-conserved phenylalanine (ALF) motif (Yeats et al., 2003); Laminin_G_3 (PF13385); LamG-like jellyroll fold domain (Liu et al., 2007; Weyer et al., 2007); HintN domain (Perler, 1998). Individual protein schemas were generated using Illustrator of Biological Sequences (Liu et al., 2015) with manual adjustment.

### Antidote characterization

HaloBlast subjects from the searches with REIS_1423 and NF27_IC00040 as queries were compiled and aligned with MUSCLE (default parameters), with the entire alignment visualized as sequence logos using WebLogo. Additionally, CDS flanking certain HaloBlast subjects (i.e. those with NCBI reference protein accession numbers) from the searches with REIS_1424 and NF27_IC00050 were evaluated for their predicted size and presence of a conserved N-terminal sequence motif (LS/ADXE/DXQXXXW) determined to be highly conserved in subjects retrieved in Blastp searches with REIS_1423 and NF27_IC00040 as queries. Finally, HMMER (Finn et al., 2011) searches using NF27_IC00040 or NF27_IC00050 were utilized to evaluate our Blastp-based identification and compilation of both toxin and antidote datasets.

### Phylogeny estimation

#### Antidote phylogeny

Selected antidotes were aligned using MUSCLE (default parameters). A phylogeny was estimated with the WAG substitution model (gamma model of rate heterogeneity) using RAxML v8.2.4 (50). Branch support was assessed with 1,000 pseudo-replications.

#### Rickettsia phylogeny

Protein sequences (*n* = 121,310) from 92 sequenced genomes were used to estimate a genus-level *Rickettsia* phylogeny. *Rickettsia* genomes were retrieved from the NCBI Assembly database (*n* = 92). The Rapid Annotation using Subsystem Technology (RAST) v 2.0 server (Aziz et al., 2008) was used to annotate three *Rickettsia* assemblies that were not previously annotated. A total of 3,707 orthologous gene families were constructed from this data set using *fastortho*, a modified version of OrthoMCL (Feris et al., 2003), at an inflation of 1.5 and a percent identity threshold of 40%. A subset of 263 single-copy families conserved across all 92 taxa was independently aligned with MUSCLE (Edgar, 2004) using default parameters, and regions of poor alignment were masked using Gblocks (Talavera and Castresana, 2007). All modified alignments were concatenated into a single data set (74,799 positions) for phylogeny estimation using RAxML v8.2.4 (Stamatakis et al., 2005), using a gamma model of rate heterogeneity and estimation of the proportion of invariable sites. Branch support was assessed with 1,000 pseudo-replications.

### DNA and RNA extraction, PCR

For *R. buchneri* analysis, ticks from New York, New Hampshire, and Pennsylvania (see **Fig. 6** for locality information) were stored in 100% ethanol at −20°C until isolation. DNA was extracted from *I. scapularis* adults and nymphs and rickettsiae infecting ISE6 cells and using the DNeasy kit (Qiagen) as per manufacturer’s protocols for cell culture and tissue extraction, respectively. Briefly, ticks were surface sterilized with 5 min washes (1% bleach, 70% ethanol, and 1xPBS), cut into quarters with a sterile scalpel blade, incubated with kit-provided digestion buffer with proteinase K at 56°C overnight, and extracted using the tissue protocol with a final elution of 50μl of molecular grade water. Rickettsiae grown in culture were collected in their host cells, and DNA extracted using the cell culture protocol with a 50μl elution with molecular grade water. For analysis of non-target rickettsiae, all bacteria were grown in cell culture prior to DNA extraction and PCR analysis. DNA was qPCR amplified using PowerUp Sybr Mastermix (Thermo) in 20μl reactions containing 400nm of each primer and 50-100ng of DNA. Primers pairs are as follows for *R. buchneri*-specific targets: Rb-1424-120-F-5’-acaggcgtaaaactagacaatct-3’ with Rb-1424-120-R-5’-aggaaatccaagcttttcaggta-3’ for the amplification of *rCRCT* and Rb-1423-116-F-5’-gcatagggtttatagcggtgc-3’ with Rb-1423-116-R-5’-ccataagtttcttcctattgtgctt-3’ for the amplification of *rCRCA*. *Rickettsia gltA* was amplified for all rickettsiae using the following primers: CSRT-F-5’-tcgcaaatgttcacggtacttt-3’ and CSRT-R-5’-tcgtgcaattctttccattgt-3’ (Stenos et al., 2005). Reactions were amplified under the following conditions: 1 cycle for 2 min at 95°C, 45 cycles at 95°C for 15 sec and 60°C for 30 sec, followed by a melt curve analysis. All primer sets were considered positive if the cycle threshold was 37 cycles or less. All primer sets were validated for range and efficiency of amplification using pCR4-TOPO plasmid standard curves with ligated amplicons. Primer sets described in this manuscript only amplify their intended products as verified by sanger sequencing and melt curve analyses of each reaction. For visualization of qPCR products (**Fig. 6**) gel electrophoresis was performed using a 2% agarose gel with ethidium bromide straining and visualization using a gel imaging station. For transcriptional analysis, *R. buchneri* growing during log growth in ISE6 cells were collected in 600μl TRIzol (Invitrogen) and RNA extracted using the DirectZol (Zymo) kit using manufacturer’s instructions and on-column DNase treatment. RNA was further DNase treated using the RQ1 DNase (Promega) prior to enzyme removal and concentration with the Zymo Clean and Concentrator-5 kit. The iScript Select cDNA synthesis kit (Bio-Rad) was used for cDNA synthesis using random hexamers in 20μl reactions with 200ng of DNase-free RNA. RNA was determined to be free of DNA with no reverse transcriptase reactions with 200ng of RNA which resulted in no detectable DNA by qPCR. cDNA was analyzed for transcription of *rCRCT*, *rCRCA* and *I*. *scapularis* β-actin (primers IsActin-95-F-5’-aatcggcaacgagaggttcc-3’ and IsActin-95-R-5’-agttgtacgtggtctcgtgg-3’) using the same qPCR parameters as above.

## Results and Discussion

### Identifying an *R*. *buchneri*-specific gene

Our prior analysis of the first sequenced *R*. *buchneri* genome (Wikel *I*. *scapularis* colony) indicated 32% of CDS were absent from other *Rickettsia* genomes (Gillespie et al., 2012a). Given that dozens of new *Rickettsia* genomes have been sequenced and assembled since 2012, we revisited this list and allowed for candidate genes to also be present in the *R*. *buchneri* str. ISO7 assembly (Kurtti et al., 2015). Further, to increase the likelihood of a stable gene present in all *R*. *buchneri* populations, we excluded genes 1) encoded on plasmids, 2) flanked by transposases, 3) containing annotations reflecting an association with MGEs, and 4) containing Blastp profiles indicating pseudogenization (e.g., gene fragments, split genes, etc.). These collective constraints yielded a small list of candidate genes, one of which (*REIS_1424*, NCBI accession no. EER22217) was selected for further analysis.

*REIS_1424* encodes a hypothetical protein of 263 aa; the homolog in the *R*. *buchneri* str. ISO7 assembly (KDO03356) is identical yet has a different predicted start site that adds 35 aa to the N-terminus. The only remaining significant Blastp matches are from *R*. *tamurae* str. AT-1, with two CDS spanning the entirety REIS_1424 indicating a split gene (WP_215426163 and WP_032138795). This is consistent with the close phylogenetic position of *R*. *tamurae* and *R*. *buchneri* (Hagen et al., 2018); however, since *R*. *tamurae* has not been reported from the Western hemisphere, these CDS are not a concern for utilization of *REIS_1424* as a diagnostic for *R*. *buchneri* infection.

### REIS_1424 carries a cryptic toxin domain

REIS_1424-based Blastp searches outside of the *Rickettsia* taxon database yielded only two significant matches: a 2192 aa protein (hypothetical protein NF27_IC00050, KIE04387) from a rickettsial amoeba-associated endosymbiont, “*Candidatus* Jidaibacter acanthamoeba” str. UWC36 (Rickettsiales: Midichloriaceae) (Schulz et al., 2016), and a 97 aa protein (hypothetical protein E1266_17330, TDB94289) from the actinobacterium *Actinomadura* sp. 7K534 (Streptosporangiales; Thermomonosporaceae). Both alignments indicated that **1)** REIS_1424, NF27_IC00050, and E1266_17330 share over a dozen conserved residues, **2)** the REIS_1424 and NF27_IC00050 match aligns the proximal C-terminal sequences of both proteins, and **3)** E1266_17330 is truncated and lacks N-terminal sequence outside of the matches. Unlike REIS_1424, NF27_IC00050- or E1266_17330-based Blastp searches yielded many significant matches to diverse bacteria (discussed further below). However, no functional domains for the region shared across these proteins could be predicted with searches against the NCBI Conserved Domains Database or using SMART.

Given that the REIS_1424-NF27_IC00050 match spanned greater sequence in each protein (∼137 aa) and “*Cand.* J. acanthamoeba” is another rickettsial taxon, NF27_IC00050 (aa residues 2068-2192) was used as a proxy for further Blastp searches and *in silico* characterization. NF27_IC00050-based HaloBlast analysis revealed strongest similarity to certain non-proteobacterial proteins (**Fig. 2A**). All obtained sequences matched NF27_IC00050_2068-2192_ at their proximal C-termini (**Fig. 2B**). Intriguingly, this cohort of proteins (n = 155) varied greatly in size across regions outside of the conserved C-terminal sequence (**Fig. 2C**). A wide assortment of domains was predicted for these proteins, with many having modular architectures and other characteristics of **contact dependent growth inhibition** (**CDI**) and/or **Recombination hotspot** (**Rhs**) toxins (**Fig. 2D**). While no functional domain could be predicted for any of the analogous C-terminal regions, sufficient conservation was found to strongly indicate a unifying functional role (**Fig. 2E**, **Fig. S1A**). We hereafter refer to these analogous regions as **CDI-like/Rhs-like CT toxin** (**CRCT**) domains, and to larger proteins possessing them as **CRCT domain-containing proteins** (**CRCT-DCP**).

**FIGURE 2.**
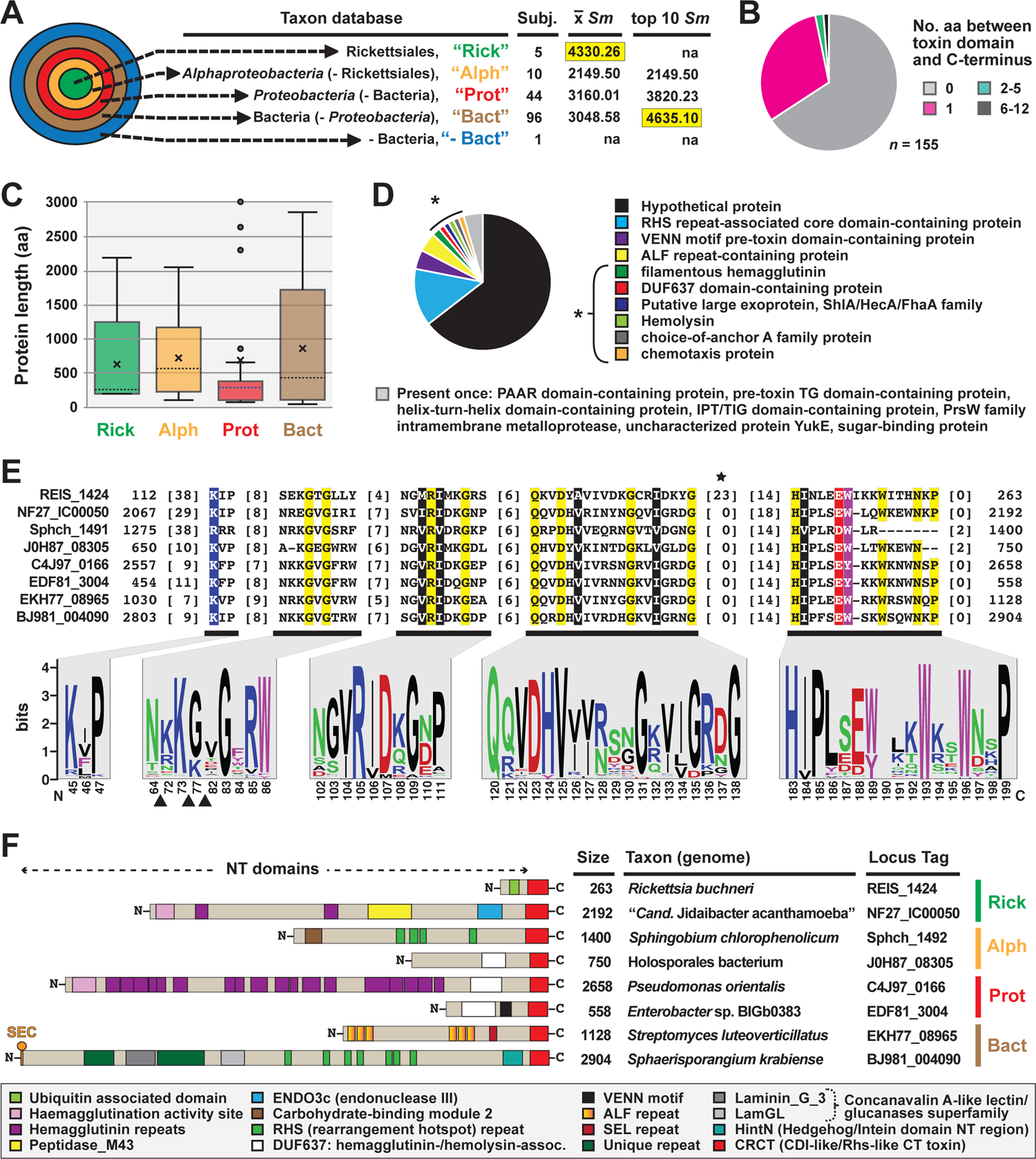
REIS_1424 of *Rickettsia buchneri* contains a C-terminal toxin domain characteristic of some bacterial contact-dependent growth inhibition (CDI) and rearrangement hotspot (Rhs) toxins. This domain was named CDI-like/Rhs-like C-terminal toxin (CRCT). (A) HaloBlast analysis for “*Candidatus* Jidaibacter acanthamoeba” CRCT of NF27_IC00050 (NCBI acc. no. KIE04387; aa residues 2068-2192). Concentric halos depict hierarchical taxonomic databases increasing in divergence from the center. Average *Sm* score (see text for details) for all subjects and top ten subjects are provided, with highest score per database highlighted. ‘na’, not applicable. All corresponding information for proteins from HaloBLAST analyses are provided in Table S1. (B-E) For compiled bacterial CRCT-containing proteins (*n* = 155): (B) number of aa residues between CRCT and C-terminus, (C) lengths for associated N-terminal regions parsed by taxonomic group (the two largest proteins for *Proteobacteria* (AZE30872, *Pseudomonas chlororaphis* subsp. Aureofaciens and TVR95235, Wenzhouxiangellaceae bacterium are not shown), (D) NCBI protein annotations, and (E) conservation within the CRCT with sequence logo illustrating alignment of 155 CRCTs (see full alignment in Fig. S1) and amino acid coloring as follows: black, hydrophobic; red, negatively charged; green, hydrophilic; purple, aromatic; blue, positively charged; star depicts unique *Rickettsia* insertion and triangles other insertions). (F) Diverse CRCT-containing proteins from select genomes.

The fusion of small**, toxin-antidote** (**TA**) pairs to the C-termini of CDI and Rhs toxins has previously been described and is thought to expand the diversity of toxic activities deployed by both CDI and Rhs systems (Aoki et al., 2010; Poole et al., 2011; Zhang et al., 2011; Ruhe et al., 2018). The extreme polymorphic nature of these TA modules indicates bacterial arms races, with selection operating on species- and strain-level recognition that shapes communities (Willett et al., 2015). For instance, many of the CRCT domain-containing proteins we identified in diverse bacteria lack the CRCT domain in closely related strains (data not shown). Furthermore, we found many small proteins containing only the CRCT domain (**Fig. 2C**, **Table S1**), as well as larger proteins carrying truncated CRCT domains (data not shown), suggesting the mobile nature of these toxins and a rapid “birth and death” process. These observations, combined with small size and limited sequence conservation, collectively challenge computational approaches for identifying these polymorphic toxins. This is evinced by HaloBlast profiles for REIS_1424 that mirror those for NF27_IC00050_2068-2192_ once a *Rickettsia*-specific insertion is removed from the query in Blastp and HMMR searches (**Fig. 2E**, **Fig. S1B**).

### REIS_1423 is a cryptic immunity antidote to REIS_1424

As CRCTs are often found as TA modules, we interrogated genes up- and downstream of *REIS_1424*, *NF27_IC00050*, and the genes encoding the 153 other identified CRCT-DCPs. This revealed probable antidotes, hereafter named **CDI-like/Rhs-like C-terminal toxin antidotes** (**CRCA**), adjacent to 37% of the 155 CRCT-DCPs (**Fig. 3A**).

**FIGURE 3.**
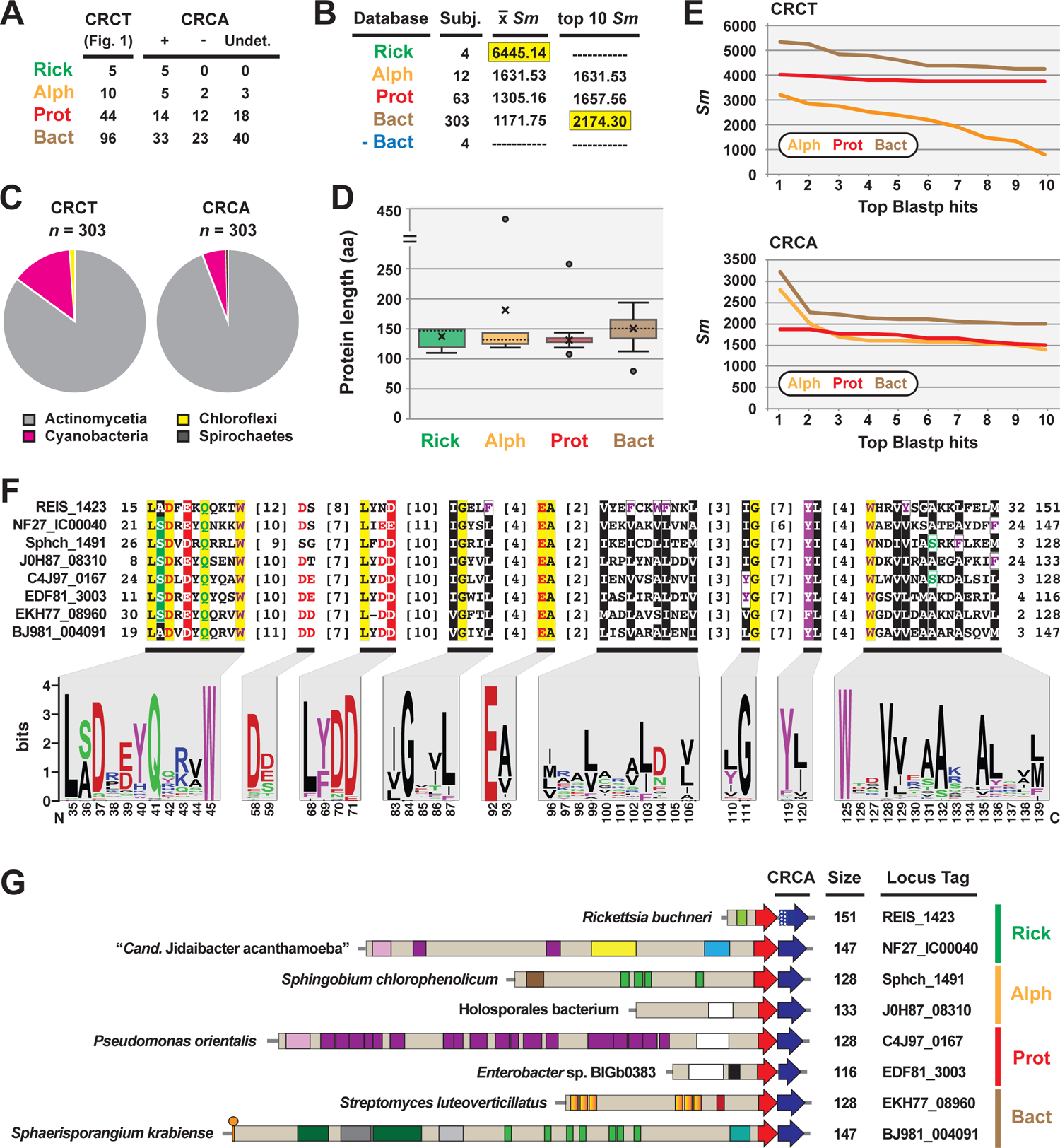
REIS_1423 of *Rickettsia buchneri* is a predicted antidote to REIS_1424. REIS_1423 and related proteins were named CDI-like/Rhs-like immunity antidotes (CRIA). (**A**) Blastp searches with REIS_1423 and NF27_IC00040 unearthed putative CRIAs adjacent to 37% of the 155 CRCT domain-containing toxins. In 61 cases, assignment of CRIAs to adjacent CRCT domain-containing toxins could not be made due to the lack of strain-specific NCBI reference protein accession numbers (non-redundant protein record (WP_) only). (**B**) HaloBlast analysis for “*Candidatus* Jidaibacter acanthamoeba” CRCT of NF27_IC00040 (NCBI acc. no. KIE04386). Concentric halos depict hierarchical taxonomic databases increasing in divergence from the center. Average *Sm* score (see text for details) for all subjects and top ten subjects are provided, with highest score per database highlighted. ‘na’, not applicable. All corresponding information for proteins from HaloBLAST analyses are provided in **Table S2**. (**C**) Taxonomic breakdown of non-proteobacterial hits retrieved in HaloBlast analysis of CRCT domain-containing toxins and CRIAs. (**D**) Lengths for CRIAs. (**E**) Top ten blastp subjects by *Sm* score (see text for details) from ‘Alph’, ‘Prot’, and ‘Bact’ searches for CRCT domain-containing toxins and CRIAs. (**F**) CRIA conservation; sequence logos illustrate alignment of 380 CRIAs (see full alignment in **Fig. S2**), with amino acid coloring as follows: black, hydrophobic; red, negatively charged; green, hydrophilic; purple, aromatic; blue, positively charged; star depicts unique *Rickettsia* insertion and triangles other insertions). (**G**) Diverse CRCT/CRIA modules from select genomes. White stipples on REIS_1423 indicate an adjusted start site.

NF27_IC00040-based HaloBlast analysis mirrored that for NF27_IC00050_2068-2192_, revealing strongest similarity to non-proteobacterial proteins (**Fig. 3B**). Taxonomic breakdown of these non-proteobacterial proteins for both CRCT-DCPs and CRCAs revealed a majority from Actinomycetia and Cyanobacteria genomes (**Fig. 3C**). While all CRCAs are strongly constrained in length (∼140 aa), with only a few proteins fused to partial genes with unrelated functions (**Fig. 3D**), Blastp profiles for the best scoring matches parsed by taxonomy strongly indicate ***Rickettsia* CRCT/CRCA** (**rCRCT/CRCA**) modules were acquired from distant non-proteobacteria via LGT (**Fig. 3E**).

Like analyzed CRCTs, predicted CRCAs possess enough conservation to indicate a common function (**Fig. 3F**, **Fig. S2**). Still, efforts to thread either CRCTs or CRIAs to solved structures of CdiA-CT/CdiI toxin/immunity complexes (Morse et al., 2012; Beck et al., 2014; Johnson et al., 2016; Michalska et al., 2018) were futile. Many of the characterized CdiA-CT toxins are from proteobacterial species and function as Rnases, specifically targeting tRNAs or rRNAs (Willett et al., 2015). Using SMART or searching against the NCBI CDD did not indicate CRCTs harbor nuclease activities, and no similarity of CRCAs to CdiI domains (e.g., cd20694: CdiI_Ct-like) or members of the SUKH superfamily of immunity proteins (Zhang et al., 2011) could be made. CdiI antidotes have been hypothesized to drive CdiA-CT/CdiI module diversification since they evolve faster than CdiA-CT toxins (Nikolakakis et al., 2012). While we observed greater conservation in CRCTs (**Fig. 2E**) versus CRCAs (**Fig. 3F**), HaloBlast and HMMER (data not shown) searches recovered more CRCA (**Fig. 3B**) versus CRCT-DCPs (**Fig. 2A**). Still, the presence of CRCT/CRCA modules across diverse bacterial phyla, with some drastic differences in lifestyle (i.e. obligate intracellular versus extracellular, eukaryote-dependent versus environmental, etc.) indicates a common universal cellular target of CRCTs such as membrane, DNA, or RNA previously characterized for all other studied CdiA-CT/CdiI modules (Aoki et al., 2009, 2010).

### A rickettsial CRCT/CRCA module mobilized to a eukaryote?

We identified a eukaryotic genome harboring a possible LGT of a rickettsial CRCT-DCP/CRCA system. The genome of the smooth cauliflower coral (*Stylophora pistillata*) contains a gene encoding a large Rhs-like toxin (YbeQ) that was assembled on an unincorporated scaffold with other bacterial-like genes (**Fig. 4**). YebQ has highest similarity to a few smaller rickettsial proteins that are remnants of degraded Rhs-like toxins (see next section), yet consistent similarity to larger toxins from non-proteobacterial genomes (**Table S3**). While YebQ does not contain a CRCT domain, smaller *S*. *pistillata* genes were found encoding the CRCT and CRCA domains with higher similarity to rickettsial equivalents than other bacteria (**Table S3**). This attests to the mobile nature of CRCT/CRCA modules and their tendency to incorporate into larger bacterial toxins. It also resonates on our prior work showing another aquatic animal, the placozoan *Trichoplax adhaerens*, contains LGTs from bacteria (particularly rickettsiae) (Driscoll et al., 2013). While the presence of introns in many of these bacterial-like *S*. *pistillata* gene models supports integration, mis-assembly of reads from a rickettsial endosymbiont is also possible.

**FIGURE 4.**
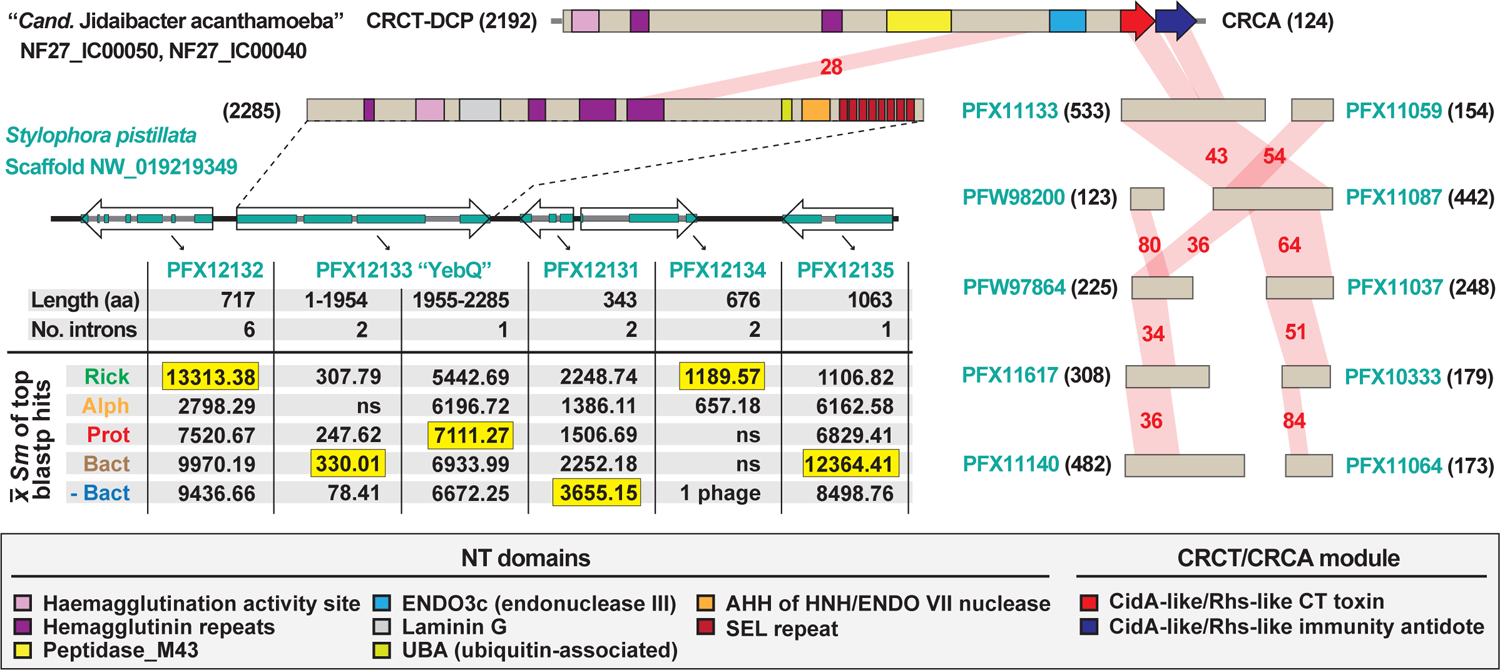
The mobile nature of CRCT/CRCA modules captured in a eukaryotic genome assembly. In blastp searches against the NCBI “non-bacteria” database, the CRCT/CRIA module of “*Candidatus* Jidaibacter acanthamoeba” str. UWC36 consistently hit only predicted proteins from the smooth cauliflower coral (*Stylophora pistillata*) genome. *Left*, the large N-terminal region of NF27_IC00050 is similar to a large protein (PFX12133, 2285 aa) encoded on a five-gene *S. pistillata* scaffold (NW_019219349). PFX12133 domain architecture (descriptions in gray inset at bottom) is reminiscent of large, multi-domain hemagglutinin-like RHS toxins that may or may not carry CRCT domains (see Fig. 2 and Fig. 3). The HaloBlast profile of PFX12133 and adjacent proteins indicates either rampant bacterial gene incorporation into the *S. pistillata* genome or mis-assembly of bacterial sequencing reads from *S. pistillata*-associated microbes. See text and Fig. 2 legend for description of HaloBlast. PFX12132: stomatin-like protein 2, mitochondrial; PFX12133: uncharacterized protein YbeQ; PFX12133: uncharacterized protein YqaJ; PFX12133: hypothetical protein AWC38_SpisGene23959; PFX12135: DNA polymerase I. *Right*, the complete or partial CRCT/CRCA module was detected in ten smaller predicted *S. pistillata* proteins encoded by genes on scaffolds not incorporated into the *S. pistillata* assembly.

### Diverse CRIA/CRCT modules are recurrent in *Rickettsia* genomes

Inspection of the genomic region where rCRCT/CRCA modules have inserted revealed two interesting findings, both of which further attest to the mobile nature of these polymorphic TA modules and their rapid birth and death process. First, the rCRCT/CRCA loci of *R*. *buchneri* and *R*. *tamurae* occur in a recombination hotspot adjacent to the SecA gene (**Fig. 5A**). This region is highly variable across *Rickettsia* genomes (**Fig. S3A**) and contains small CDS with matches to the *S*. *pistillata* YebQ Rhs toxin described above (**Fig. S4A**). Despite extraordinary variability in the number and size of CDS in this region across *Rickettsia* genomes, a conserved tRNA-Ala^TGC^ locus is always present, corroborating prior observations for CidA-CT/CdiAI modules often inserting near tRNA genes in bacterial genomes.

**FIGURE 5.**
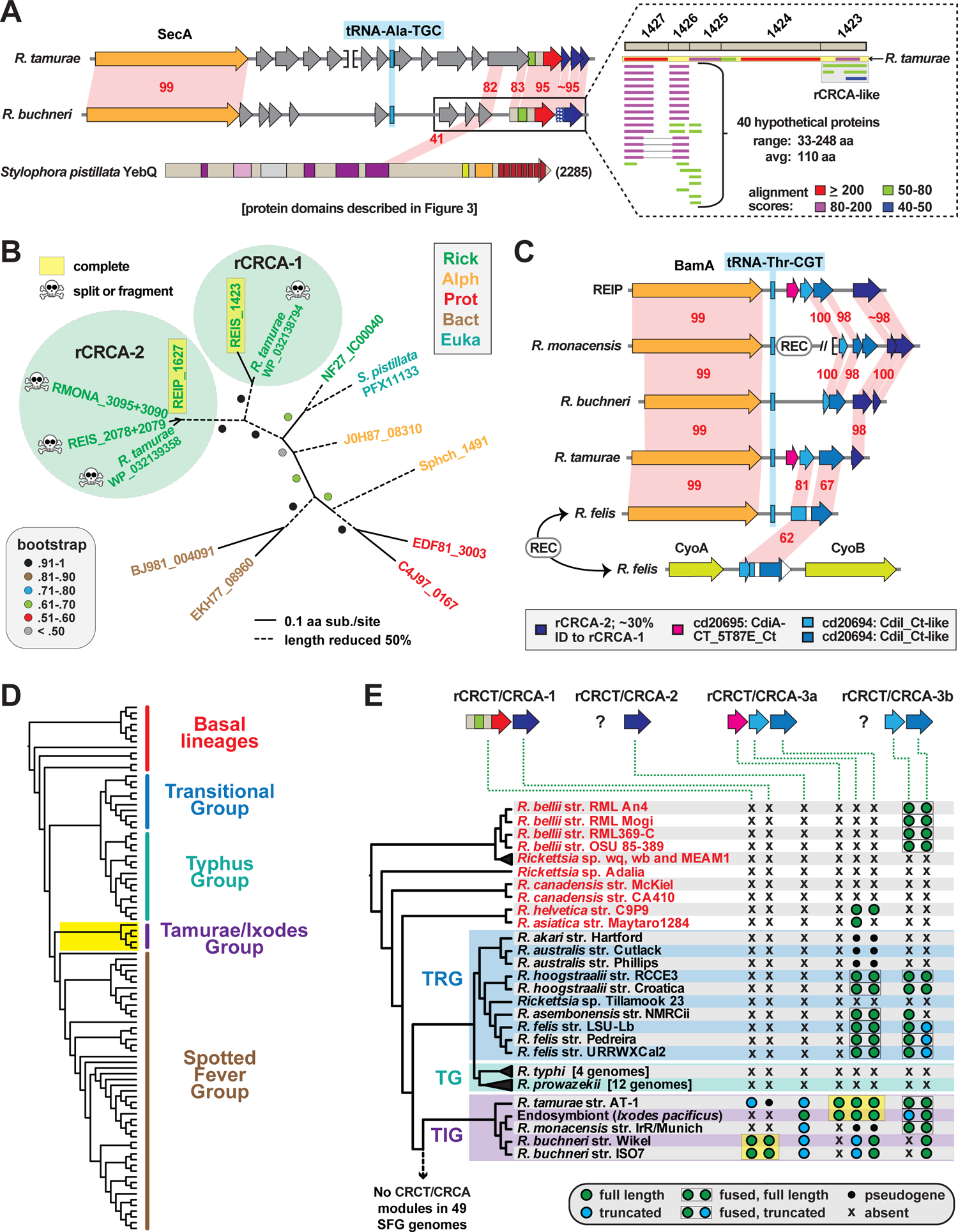
Evolution of *Rickettsia* CRCT/CRCA modules. (**A**) Comparison of *R*. *tamurae* and *R*. *buchneri* genomic regions containing *Rickettsia* CRCT/CRCA module 1 (rCRCT/CRCA-1). Gray genes encode hypothetical proteins. Limited similarity between *R*. *buchneri* REIS_1425 and *Stylophora pistillata* YebQ is shown. Dashed inset: subjects retrieved from a Blastp search against the NCBI nr database using five concatenated *R*. *buchneri* proteins (REIS_1427-REIS_1423) as the query (further information provided in **Fig. S4A**). Gray box illustrates additional rCRCA-like sequences identified by this search. (**B**) rCRCA-2 genes are divergent from rCRCA-1 and pseudogenized in all Tamurae/Ixodes Group (TIG) rickettsiae except *Rickettsia* endosymbiont of *Ixodes pacificus*. Phylogeny of rCRCA-1, rCRCA-2 and other CRCA proteins was estimated with the WAG substitution model (gamma model of rate heterogeneity) using RAxML v8.2.4 (Stamatakis, 2014). Branch support was assessed with 1,000 pseudo-replications. Final ML optimization likelihood was −3490.9. (**C**) rCRCT/CRCA modules 2 and 3a are mostly clustered near the BamA gene in some *Rickettsia* genomes, while rCRCA-3b genes occur between genes encoding the cytochrome oxidase subunits 2 (CyoA) and 1 (CyoB) in certain *Rickettsia* genomes (see **Fig. S3** for illustration of genomic regions near *secA*, *bamA* and *cyoA*/*B* loci). (**D**) *Rickettsia* phylogeny estimated from 92 genomes (see **Fig. S5** for more information). Yellow highlighting depicts TIG rickettsiae as the only species harboring complete CRCT/CRCA modules. (**E**) Phylogenomics analysis of rCRCT/CRCA modules. Yellow highlighting indicates complete rCRCT/CRCA modules. Further information is provided for rCRCT-1 (**Table S1**), rCRCA-1 (**Table S2**), rCRCA-2 (**Fig. S4B**), rCRCA-3a and rCRCA-3b (**Fig. S4C**), and rCRCT-3a (**Fig. S4D-G**).

Second, Blastp searches with a concatenated query (REIS_1427-REIS_1423) revealed matches to additional rCRCA proteins, indicating other rCRCT/CRCA modules elsewhere in *Rickettsia* genomes (**Fig. 5A**, dashed box). We designated these divergent rCRCA proteins as components of “rCRCT/CRCA-2” modules, with the above-described system named “rCRCT/CRCA-1” modules. Like the rCRCA-1 protein of *R*. *tamurae*, most rCRCA-2 genes are truncated or fragmented, yet a complete protein was found for the *Rickettsia* endosymbiont of *Ixodes pacificus* (hereafter REIP) (**Fig. S4B**). Despite limited similarity (∼30 %ID), an estimated phylogeny grouped rCRCA-1 and rCRCA-2 proteins together to the exclusion of other CRCA proteins (**Fig. 5B**; **Fig. S4B**). *rCRCA-2* loci all mapped to a second recombination hotspot in *Rickettsia* genomes adjacent to the conserved BamA and tRNA-Thr^CGT^ genes (**Fig. 5C**; **Fig. S3B**). As with rCRCA-1, we could not identify similarity between rCRCA-2 proteins and domains of CdiI proteins or SUKH immunity proteins.

Searching upstream of *rCRCA-2* for potential cognate toxins instead yielded additional genes encoding CRCA antidotes that are duplicated and highly divergent from rCRCA-1 and rCRCA-2 proteins (**Fig. 5C**). This arrangement of arrayed immunity antidotes is more characteristic of *cdiA-CT/cdiI* loci in many proteobacterial genomes (Aoki et al., 2010). Indeed, we were able to identify CdiI-like domains in these proteins using the NCBI Conserved Domains Database (cd20694). Accordingly, we named these antidotes rCRCA-3 proteins. Further inspection of rCRCA-3 genes identified eight genomes with additional copies found in a third recombination hotspot between the genes encoding the cytochrome oxidase subunits 2 (CyoA) and 1 (CyoB) (**Fig. 5C**; **Fig. S3C**). Seven *Rickettsia* genomes have rCRCA-3 genes in both the BamA and CyoA/B recombination hotspots, indicating recent recombination between these loci (**Fig. S4C**). We designated rCRCA-3a and rCRCA-3b proteins to distinguish between those located at the BamA or CyoA/B recombination hotspots, respectively. At the BamA recombination hotspot, we identified cognate rCRCT-3a toxins with CdiA-CT-like domains (cd20695) in the *R*. *tamurae* and REIP genomes (**Fig. 5C**).

These two toxins have strongest similarity to counterparts in proteobacterial genomes, particularly *Pseudomonas* and *Moraxella* species (**Fig. S4D**,**E**). We modeled the *R*. *tamurae* rCRCT-3a toxin to the CdiA-CT structure of *Cupriavidus taiwanensis* (Kryshtafovych et al., 2018) with high confidence, indicating rCRCT-3 toxins are unrelated to rCRCT-1 toxins (**Fig. S4F**,**G**). The rCRCA-3 antidotes could not be modeled to the CdiI structure of *C. taiwanensis* or any other CdiI structures, making the association of rCRCT-3 with rCRCA-3 supported by genome proximity alone.

Collectively, this analysis of rCRCT/CRCA genes in *Rickettsia* genomes illuminates recurrent genome integration, possibly in larger Rhs toxins that have degraded over time. The presence of complete, yet divergent rCRCT/CRCA modules in different species of the **Tamurae/Ixodes Group** (**TIG**) rickettsiae (**Fig. 5D,E**) indicates weaponry for interbacterial antagonism may be functional for these species and implicates a previously unrealized mechanism for rickettsial competition in the same arthropod host.

### Probing rCRCT/CRIA-1 for a role in *R*. *buchneri* biology

#### Consideration for another factor behind a putative mutualism

Factors that that distinguish parasitic rickettsiae from species exhibiting other host associations are sorely needed for Rickettsiology. Previously, we searched for genes underlying potential mutualism within the intriguing *R*. *felis* system, wherein typical strains infect blood-feeding arthropods (mostly the cat flea, *Ctenocephalides felis*) yet another has developed a tight host association with a non-blood-feeding insect (the booklouse *Liposcelis bostrychophila*) (Gillespie et al., 2015a). Only the *L. bostrychophila*-infecting strain harbored the unique plasmid, pLbAR, which we postulated encoded factors inducing parthenogenesis in booklice since sexually reproducing populations are only observed in the absence of *R*. *felis* (Yang et al., 2015).

A TA module on pLbAR was found to have similarity to gene pairs in *Wolbachia* reproductive parasites that were later characterized as the factors underpinning cytoplasmic incompatibility (or male sterilization) (Beckmann et al., 2017; LePage et al., 2017). We later reported on the widespread occurrence in intracellular species of highly diverse TA modules with rudimentary similarity to the *Wolbachia* and *R*. *felis* TA modules (Gillespie et al., 2018). Despite its high frequency of infection in *I*. *scapularis* populations, we found no evidence for these TA modules in *R*. *buchneri*, although other different factors inducing reproductive parasitism could still be present.

The presence of genes in *R*. *buchneri* that encode AGAB synthesis/resistance proteins and PKS-containing NRPS modules hint at arsenals of antibiotics and 2° metabolites possibly used for defense against certain microbes also infecting blacklegged tick. Furthermore, two copies of BOOM suggest this species supplements the blacklegged tick diet with biotin. Absence (AGAB synthesis/resistance and PKS-containing NRPS) (Hagen et al., 2018) or scarce (BOOM) (Driscoll et al., 2020) distribution of these genes in other *Rickettsia* genomes suggests they are at least utilized for functions generally not employed by other rickettsiae. For instance, a similar type I PKS of a honey bee endosymbiont has recently been shown to suppress growth of fungal pathogens and protect bee brood from infection (Miller et al., 2020). We previously showed that, unlike the conserved AGAB synthesis/resistance genes, the PKS NRPS module is variable in gene content across *R*. *buchneri* strains from different populations (Hagen et al., 2018). More recently, the PKS NRPS module and the AGAB synthesis/resistance gene array were investigated for their possible roles in limiting superinfection of pathogenic rickettsia in tick cells infected with *R*. *buchneri* str. ISO7 (Cull et al., 2022). Despite demonstrating that *R*. *buchneri* substantially reduced superinfection by pathogenic *R*. *parkeri* in cell culture, no anti-bacterial activity against either *R*. *parkeri* or extracellular bacteria (*Escherichia coli* and *Staphylococcus aureus*) was shown by the unknown product(s) of these loci, leaving their function in *R*. *buchneri* unclear.

#### rCRCT/CRCA modules are tailored for interspecific antagonism

The sequence profiles that strongly indicate rCRCT/CRCA modules were acquired from distant non-proteobacteria (**Fig. 2A**; **Fig. 3A-C,E**) are similar to those we previously reported for AGAB synthesis/resistance proteins (Gillespie et al., 2012a) and PKS-containing NRPS modules (Hagen et al., 2018). This implicates LGT in shaping these factors that likely underpin interbacterial antagonism. While the target of the product of the PKS-containing NRPS module is hard to predict, it is likely that the aminoglycoside synthesized by the AGAB gene array does not target rickettsiae, as this class of antibiotics is ineffective against *Rickettsia* species (Rolain et al., 1998) and overall generally poor for destroying intracellular bacteria (Maurin and Raoult, 2001). We find the rCRCT/CRCA-1 module a more appealing candidate for intrageneric antagonism given most described CdiA-CT/CdiI and Rhs TA modules arm bacteria for battle with self or closely related species (Aoki et al., 2010; Poole et al., 2011; Zhang et al., 2011).

While often tethered to larger N-terminal sequences with toxic activity, CdiA-CT domains themselves are toxins (Nikolakakis et al., 2012). Curiously, the N-terminal sequence of REIS_1424 contains a putative **ubiquitin-associated** (**UBA**) domain that is separated from the C-terminal CRCT domain by two “VENN” motifs, which typically demarcate the CdiA-CT domain from the remaining protein in most other CDI systems (Aoki et al., 2010; Zhang et al., 2011) (**Fig. S6**). If functional, this UBA domain may recruit host ubiquitin to the target cell and render it vulnerable to proteasomal destruction. Combined with toxin effect of the C-terminal domain, rCRCT-1 would be highly effective for targeting conspecific bacteria in the intracellular environment.

#### *rCRCT/CRIA-1* expression and possible routes for secretion

During infection of ISE6 cells, which were originally derived from *I*. *scapularis* embryos (Munderloh et al., 1999), *R*. *buchneri* expresses the REIS_1424/REIS_1423 module (**Fig. 6A**). The same primers did not amplify product in PCR reactions using DNA template from 10 diverse rickettsiae, illustrating their specificity and efficacy for future surveys of blacklegged tick populations. This is demonstrated by testing the primers on a small sampling of blacklegged ticks from three different populations throughout the *I*. *scapularis* range (**Fig. 6B**). Future sequencing of these loci will provide resolution on the conservation rCRCT/CRCA-1 and whether there is evidence for an arms race between

**FIGURE 6.**
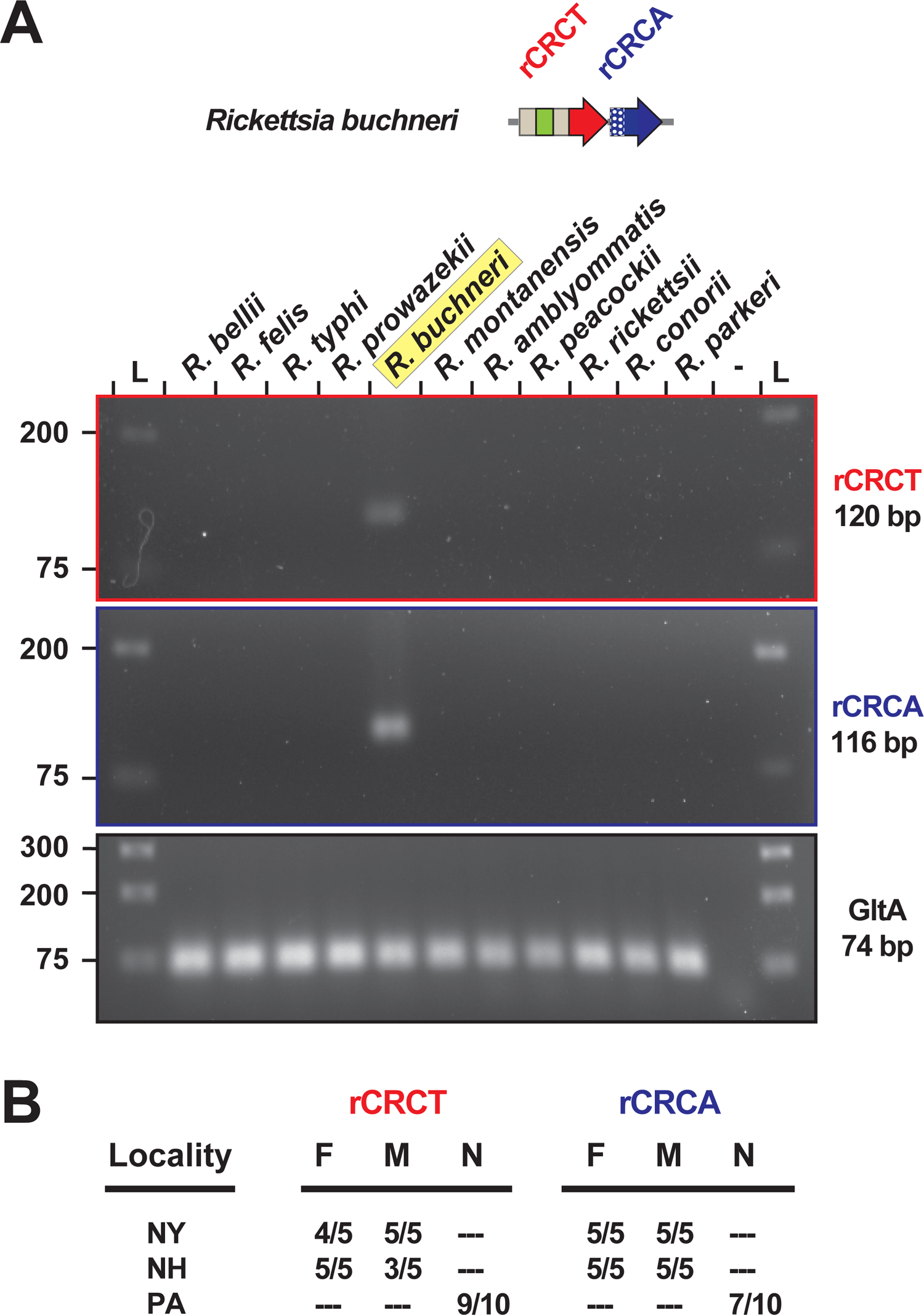
REIS_1424/REIS_1423 is a species-specific marker. (**A**) PCR assay for *rCRCT* and *rCRCA* in 11 different rickettsiae. Citrate synthase (*gltA*) was used as a positive control. (**B**) The same PCR assay was conducted on *I*. *scapularis* from three populations.

#### *R*. *buchneri* that have diverged throughout the *I. scapularis* geographic range

As *R*. *buchneri* REIS_1424 lacks a predicted signal sequence, it is likely secreted via one of two Sec-independent pathways: **type I secretion system** (**T1SS**) or **type IV secretion system** (**T4SS**), which are both conserved in rickettsiae (Gillespie et al., 2015b). Inspection of the neighborhood loci around select CRCT/CRCA modules indicates several diverse secretion pathways for some CRCT-DCPs, including T4SS (*Sphingobium chlorophenolicum*) and **type VI secretion system** (**T6SS**) (*Sphaerisporangium krabiense*), as well as typical CDI systems (*Pseudomonas orientalis* and *Enterobacter* sp. BIGb0383) with nearby *cdiB* loci, which encode the outer membrane CdiB that translocates the large CdiA protein as a **type Vb secretion system** (**T5bSS**) (**Fig. S7A**-**G**). Using CdiB as a query in Blastp searches against the major taxa harboring CRCT/CRCA modules revealed their scarcity in Rickettsiales (none detected in rickettsiae), Actinobacteria, Chloroflexi, and Spirochaetes genomes, but widespread distribution in other proteobacterial and Cyanobacteria genomes (**Fig. S7H**,**I**). This indicates that bacteria employing the CRCT/CRCA modules we describe here for warfare utilize a variety of secretions systems (e.g., T1SS, T4SS, T5bSS, T6SS, and likely others) consistent with the plethora of secretory pathways now characterized for diverse TA modules involved in interbacterial antagonism (Ruhe et al., 2018; Lin et al., 2020). The lack of *cdiB* genes and evidence that rCRCT toxins were originally appended to larger proteins indicates these modular Rhs toxins were once widespread in *Rickettsia* genomes. This is reminiscent of the large modular toxins we identified across numerous intracellular bacteria that encode a myriad of eukaryotic-like domains, some of which function in commandeering host reproduction (Gillespie et al., 2018). Like rCRCT/CRCA modules, many of these variable toxins are found adjacent to genes encoding probable antidotes, indicating a recapitulating theme for toxin architecture that persists evolutionarily and drives innovative strategies for colonizing eukaryotic hosts.

## Conclusion

Since our initial report on its genome (Gillespie et al., 2012a) and its subsequent formal species description (Kurtti et al., 2015), the appreciation for the oddity of *R*. *buchneri* relative to other rickettsiae has grown. Numerous reports on the *I*. *scapularis* microbiome now attest to the high infection rate of this microbe, particularly in females, throughout the blacklegged tick geographic range. While few reports indicate tick salivary gland infection (Steiner et al., 2008; Zolnik et al., 2016; Al-Khafaji et al., 2020), ovaries are the predominant tissue infected, consistent with the lack of reports on vertebrate infection or presence in other arthropods that co-feed on blacklegged tick hosts. This indicates a unique endosymbiosis, the intricacies of which stand to be illuminated in light of the powerful tools created by Kurtti, Munderloh and colleagues for studying this system (Munderloh et al., 1999; Simser et al., 2002; Kurtti et al., 2016; Oliver et al., 2021). Ultimately, the fitness of blacklegged ticks uninfected with *R*. *buchneri* (Oliver et al., 2021) needs to be evaluated, as does the possibility that other endosymbionts may replace *R*. *buchneri* in a mutualistic capacity.

While previously observed in ticks (Burgdorfer et al., 1981; Macaluso et al., 2002; Wright et al., 2015; Levin et al., 2018) and cell culture (Cull et al., 2022), the ability of *Rickettsia* infection to block *Rickettsia* superinfection remains a sorely understudied aspect of vector biology. Our identification and characterization of rCRCT/CRCA modules adds to a short list of factors, namely BOOM, AGAB and PKS-NRPS, that have been hypothesized to underpin a mutualism between *R*. *buchneri* and blacklegged tick. Future characterization of these factors will determine their contribution to blocking superinfection of *I*. *scapularis* by *Rickettsia* pathogens (Cull et al., 2022). When mutagenesis is someday an efficacious tool for bioengineering Rickettsiae, this line of research will offer a gene drive tool (*R*. *buchneri*) ready to disseminate into blacklegged tick populations to combat the spread of human disease agents,

## Acknowledgments

We thank Alison Luce-Fedrow (Shippensburg University), Richard Ostfeld (Cary Institute of Ecosystem Studies), and Kathryn Cottingham (Dartmouth College) for providing tick specimens, and Kevin Macaluso (University of South Alabama) and Sean Riley (University of Maryland College Park). We are grateful to Uli Munderloh and Tim Kurtti (University of Minnesota) for sharing the ISE6 cell line and *R. buchneri* strain IS07. This work was supported with funds from the National Institute of Health/National Institute of Allergy and Infectious Diseases grant R21AI156762 (JJG and TPD). TDF was supported by the Biotechnical Institute of Maryland. The content is solely the responsibility of the authors and does not necessarily represent the official views of the funding agencies. The funders had no role in study design, data collection and analysis, decision to publish, or preparation of the manuscript.

## SUPPLEMENTARY MATERIAL

**FIGURE S1.**
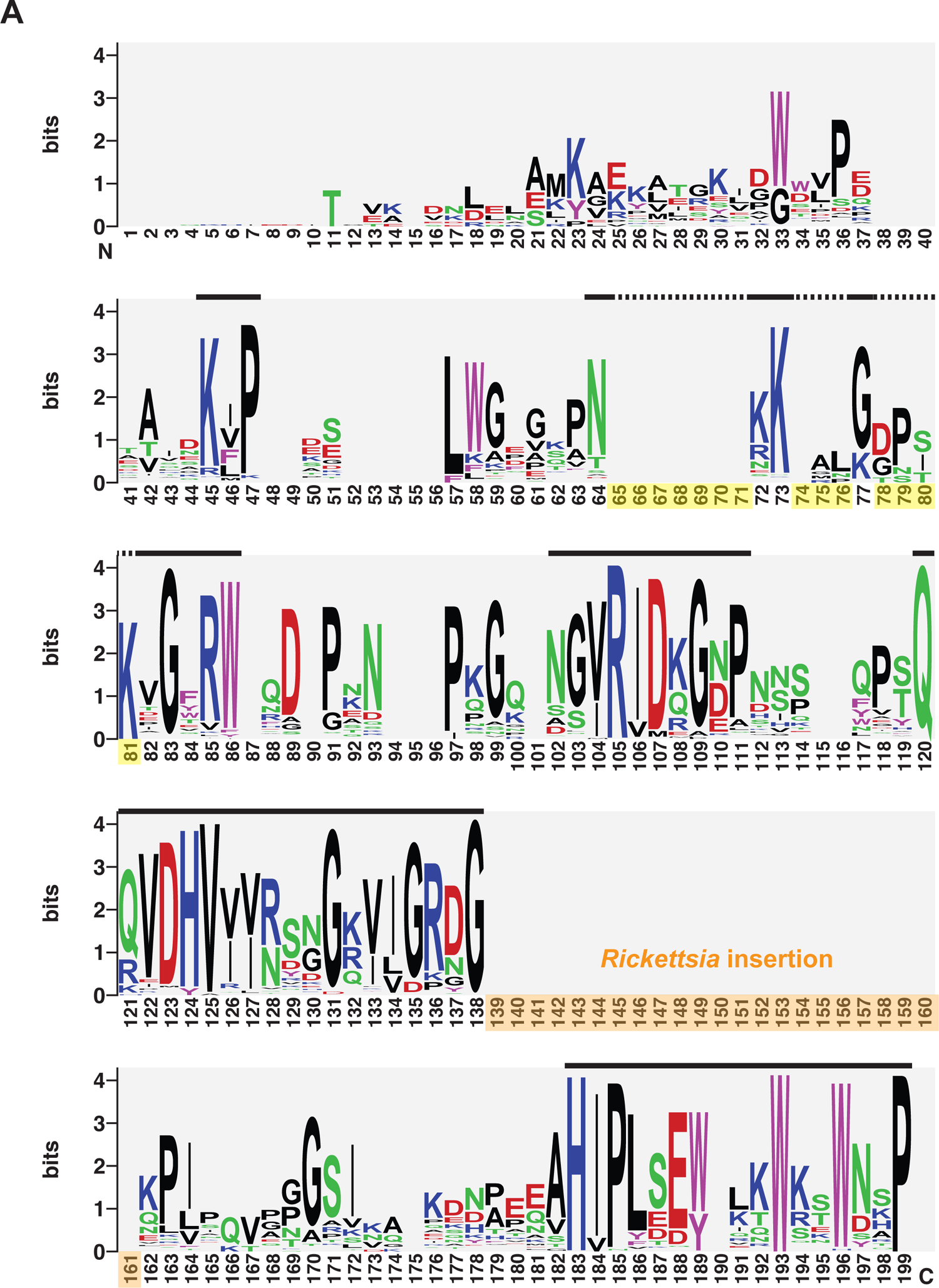
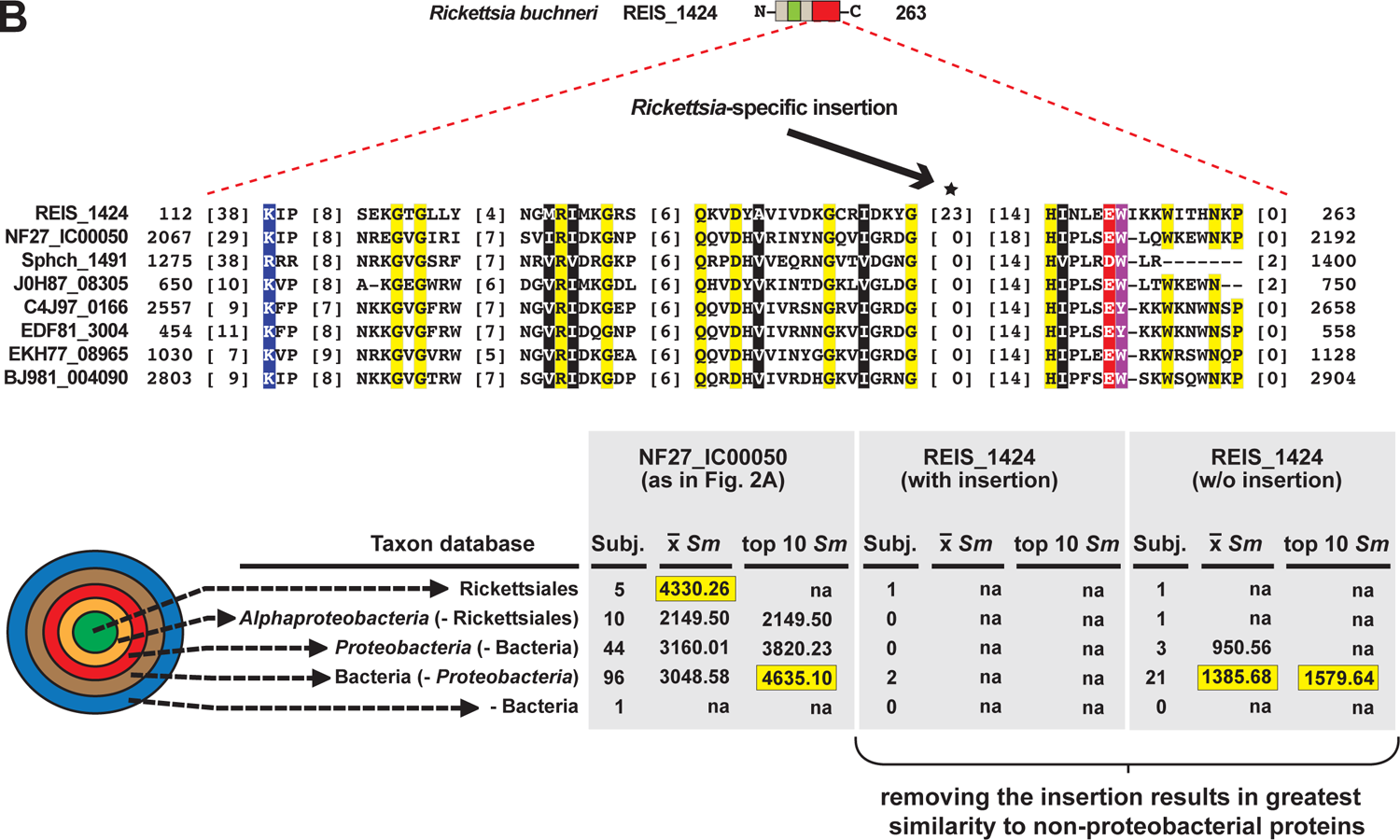
Sequence analysis of 155 predicted CRCTs integrated into diverse bacterial toxins. (**A**) Sequence logos (Crooks et al., 2004) depict complete protein alignment generated using MUSCLE, default parameters (Edgar, 2004). Information for all proteins is provided in **Table S1**. Regions shown in Figure 2 are denoted with a bar above the logos. Amino acid coloring as follows: black, hydrophobic; red, negatively charged; green, hydrophilic; purple, aromatic; blue, positively charged. Unique *Rickettsia* insertion is shown in orange. Other insertions noted by triangles in Figure 2 are highlighted yellow). (**B**) HaloBlast profiles for REIS_1424 with and without a 23 aa insertion (see Fig. 2E and text for more details).

**Table S1. Information pertaining to 155 predicted CRCTs integrated into diverse bacterial toxins.**

**FIGURE S2.**
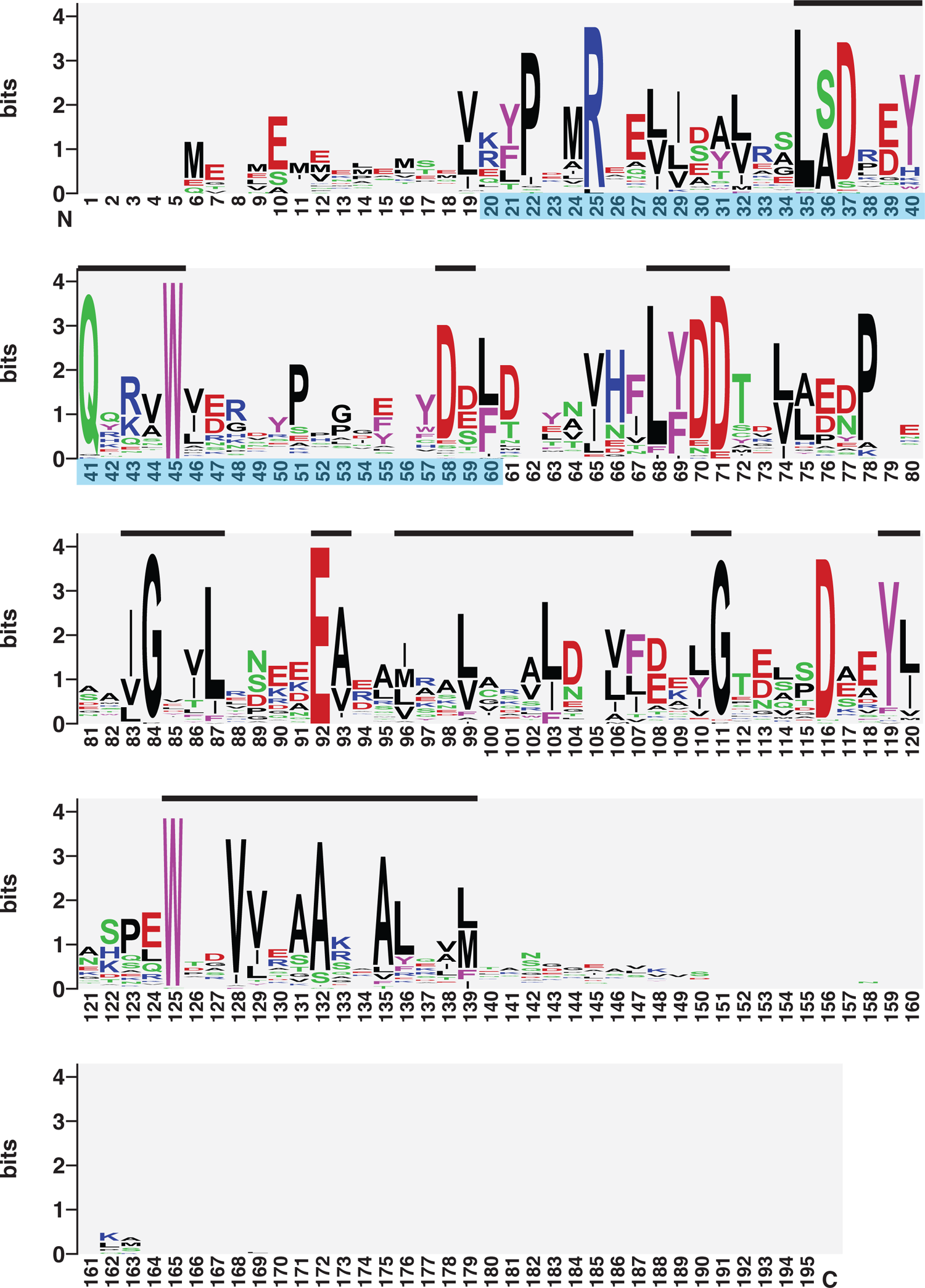
Sequence analysis of 380 predicted CRCAs. Sequence logos (Crooks et al., 2004) depict complete protein alignment generated using MUSCLE, default parameters (Edgar, 2004). Information for all proteins is provided in Table S2. Regions shown in **Figure 3** are denoted with a bar above the logos. Amino acid coloring as follows: black, hydrophobic; red, negatively charged; green, hydrophilic; purple, aromatic; blue, positively charged. Sequence spanning the adjusted start site of REIS_1423 (*Rickettsia buchneri*) insertion is shown in blue.

**Table S2. Information pertaining to 380 predicted CRCAs.**

**Table S3. Evidence for CRCT/CRCA modules in the *Stylophora pistillata* genome assembly.**

**FIGURE S3.**
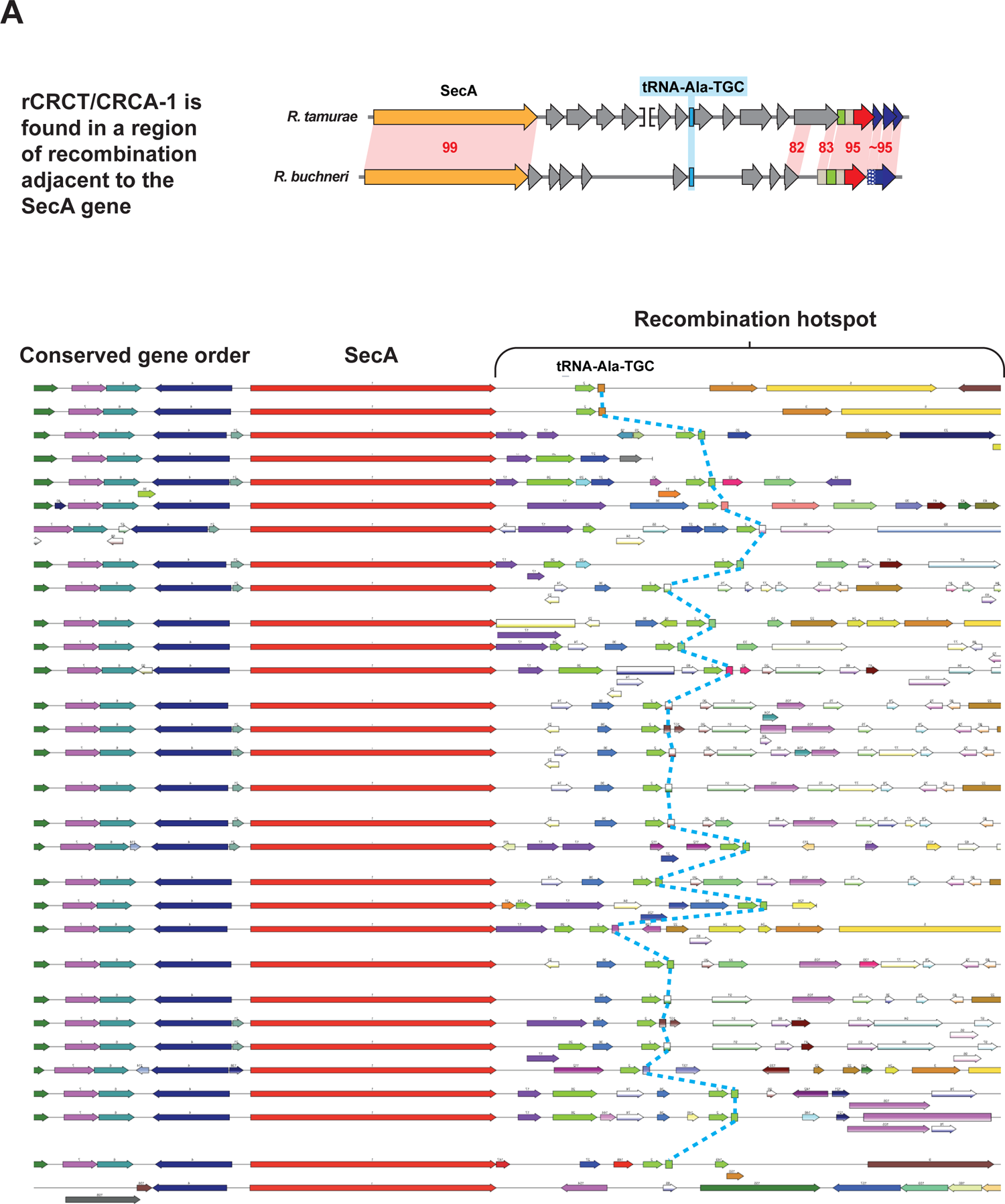
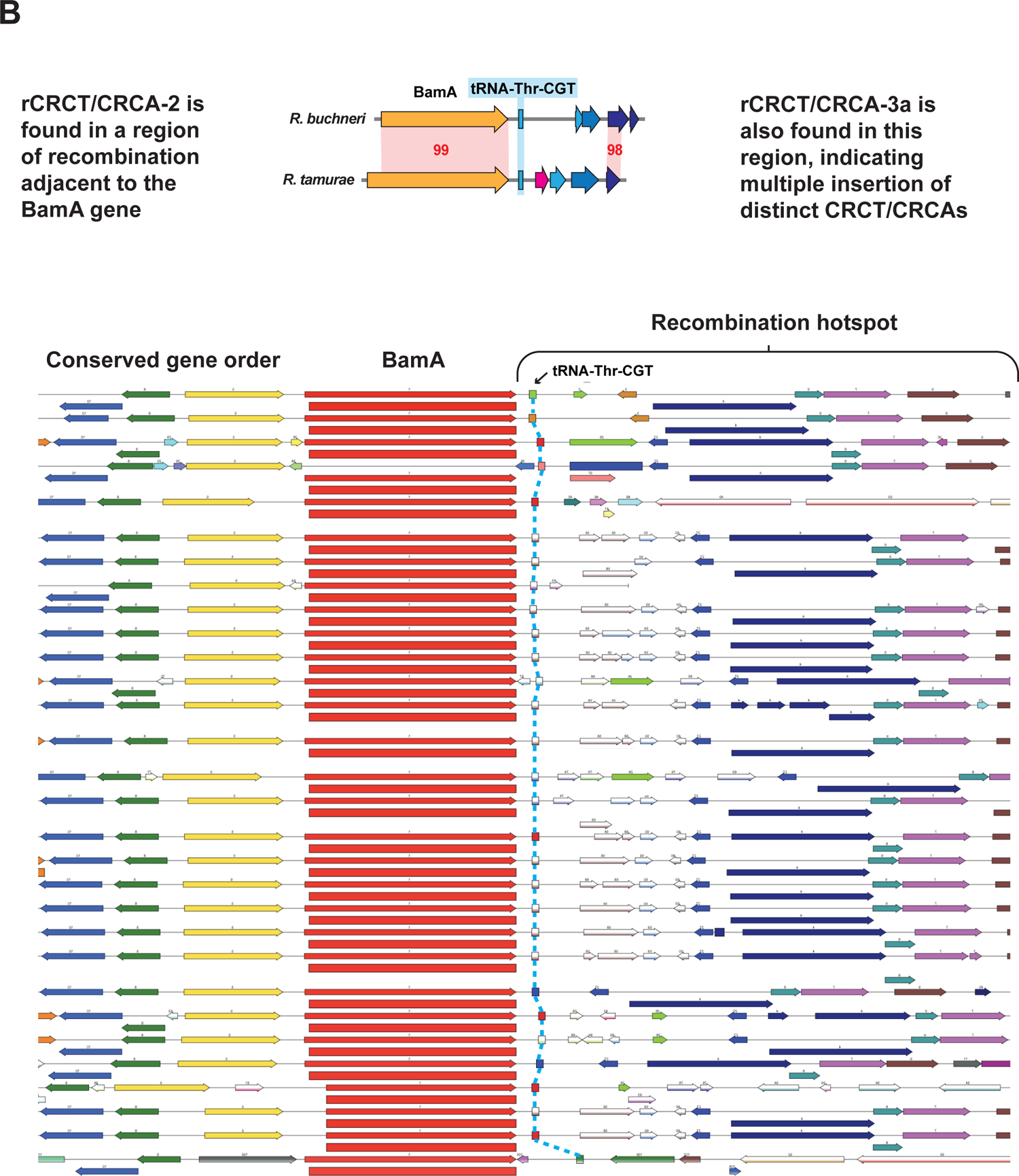
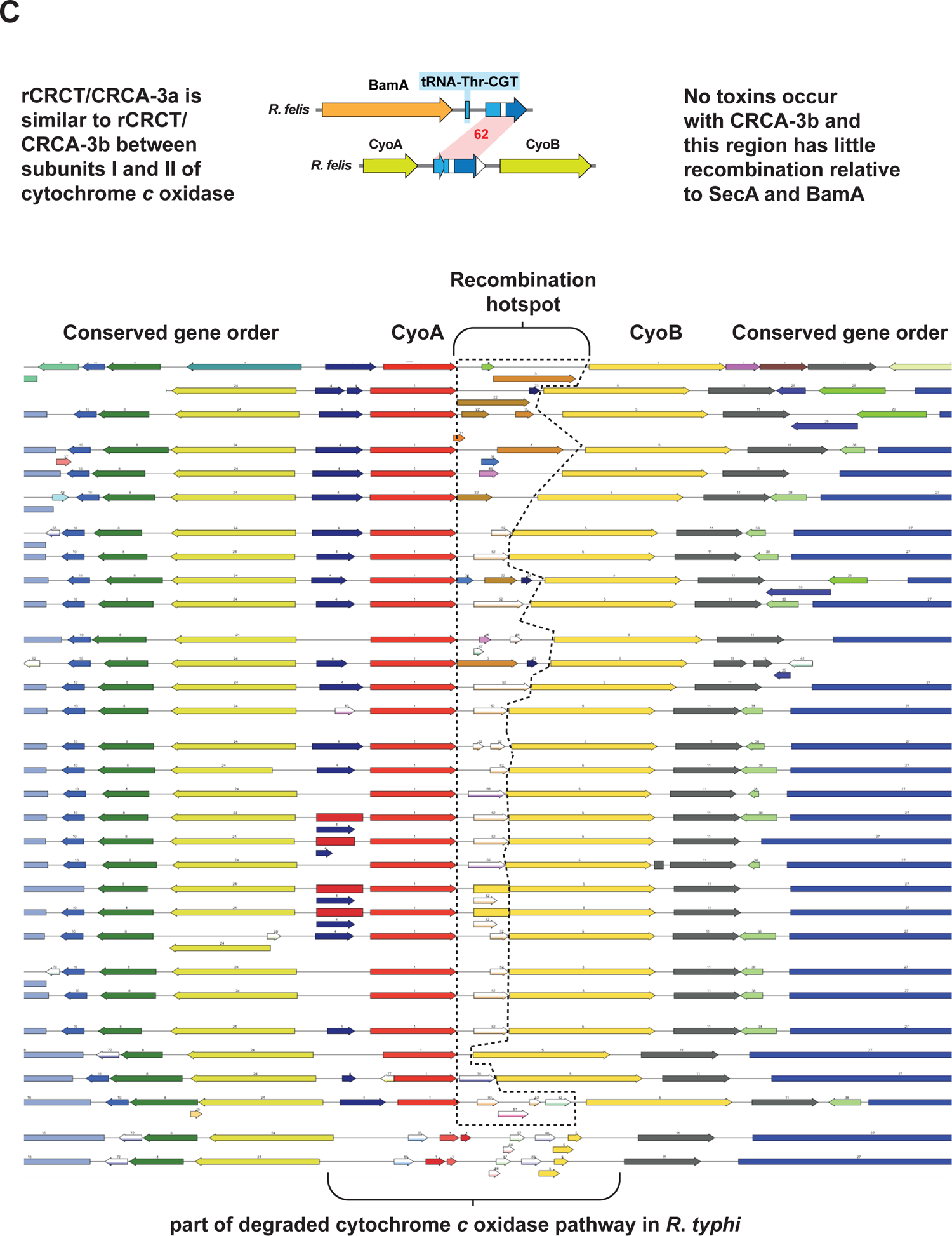
*Rickettsia* CRCT/CRCA modules occur in three recombination hotspots. (**A**) rCRCT/CRCA-1 occurs in a recombination hotspot near the SecA and tRNA-Ala^TGC^ genes. (**B**) rCRCT/CRCA-2 occurs in a recombination hotspot near the BamA and tRNA-Thr^CGT^ genes. A second TA module, rCRCT/CRCA-3a, also occurs in this region and is distinct from rCRCT/CRCA-1 and rCRCT/CRCA-2 (cd20695: CdiA-CT_5T87E_Ct, cd20694: CdiI_Ct-like). (**C**) rCRCT/CRCA-3b proteins are analogous to rCRCT/CRCA-3a and occur in a subset of *Rickettsia* genomes between *cyoB* and *cyoA*, which encode the cytochrome *c* oxidase subunits I and II, respectively.

**FIGURE S4.**
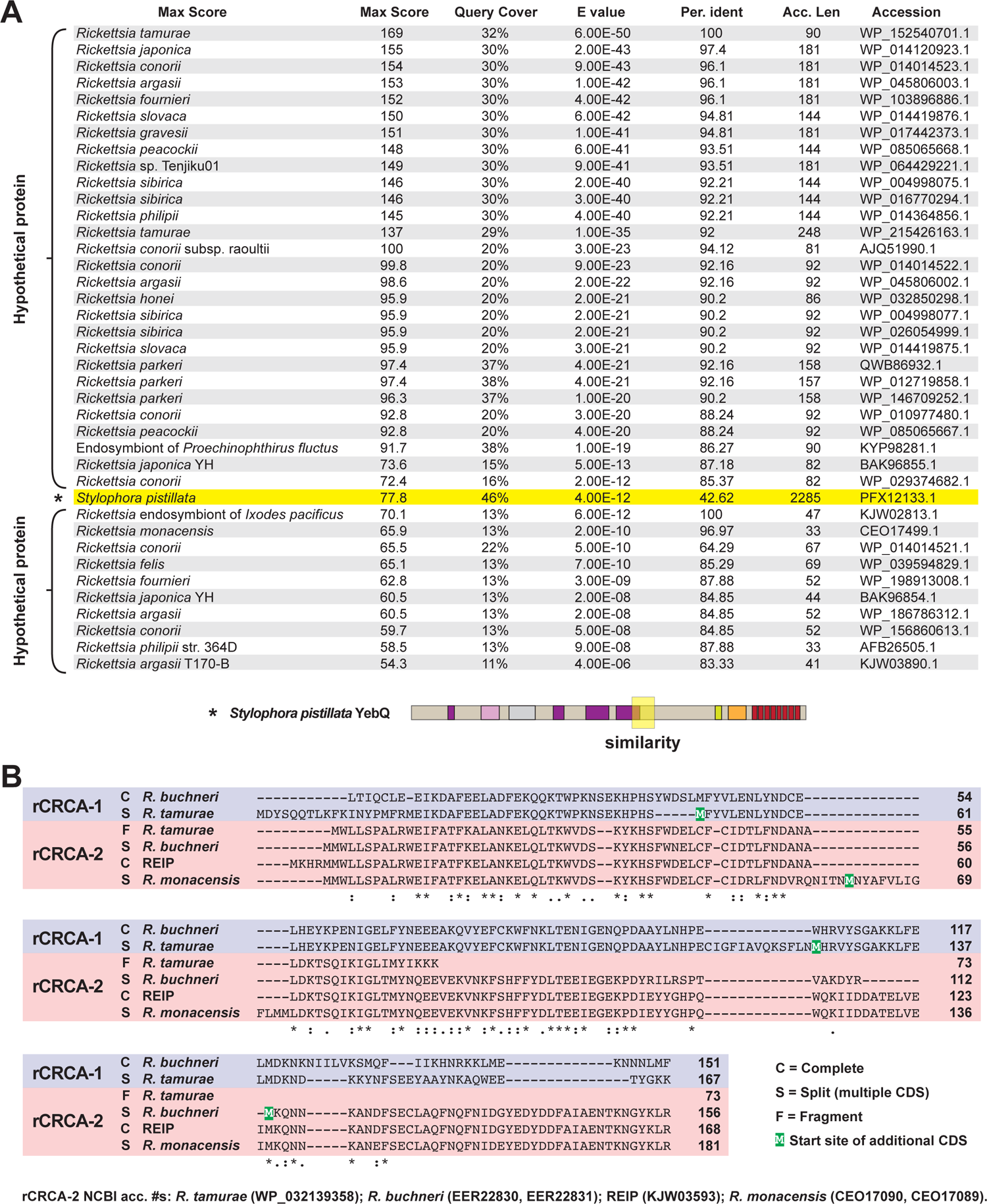
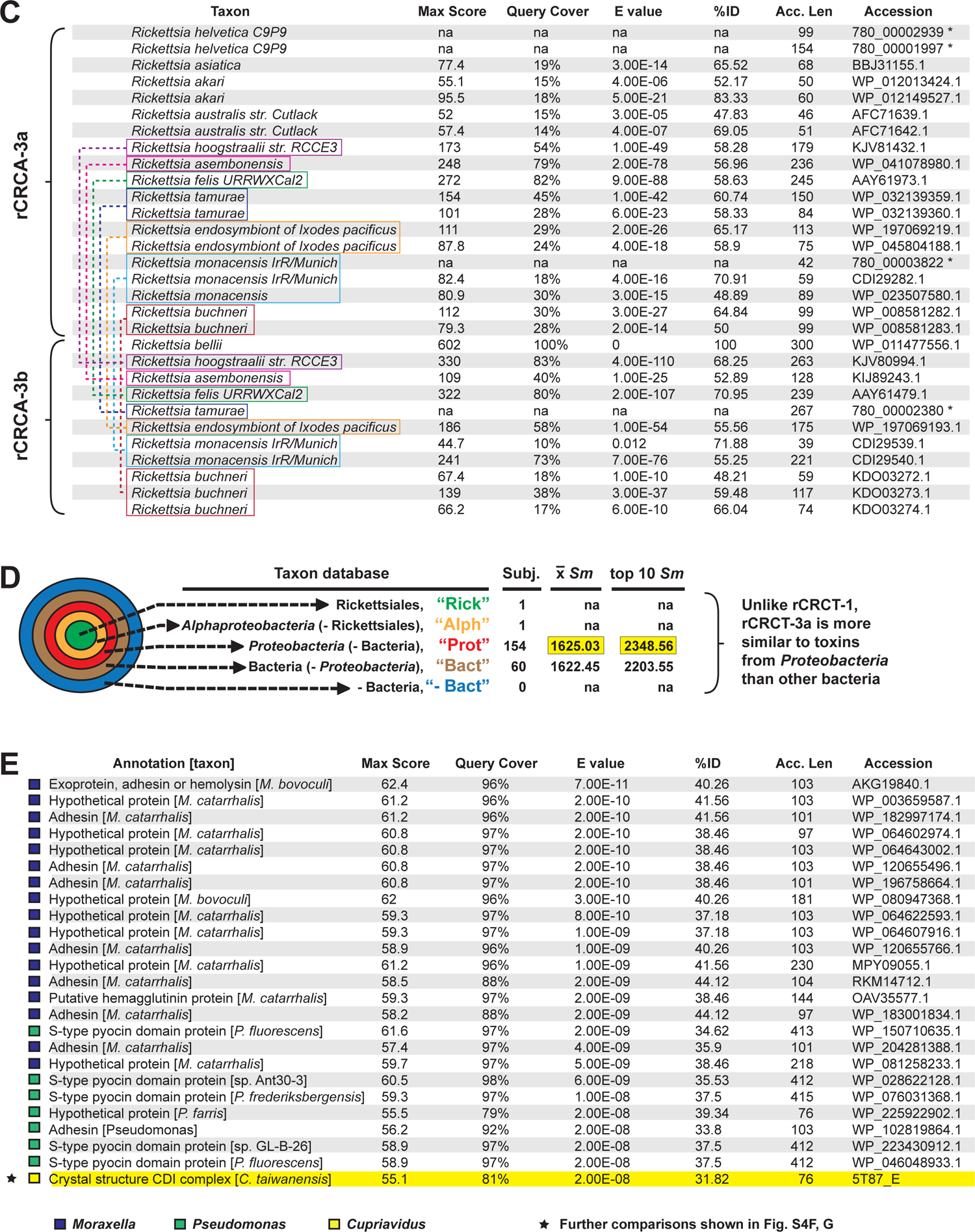
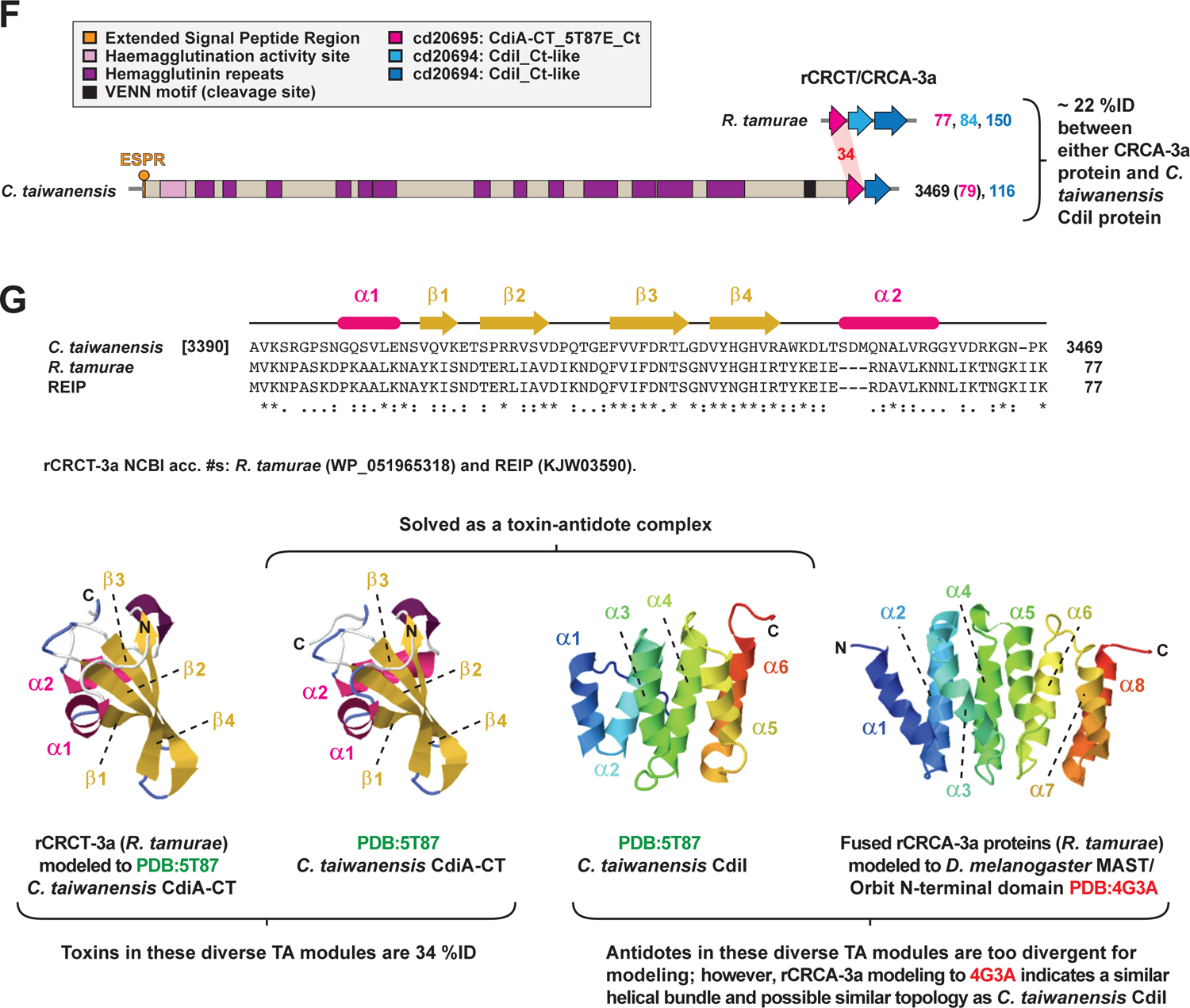
Evidence for recurrent integration of diverse CRCT/CRCA modules in *Rickettsia* genomes at recombination hotspots. (**A**) Subjects (n = 40) retrieved from a Blastp search against the NCBI nr database using five concatenated *R*. *buchneri* proteins (REIS_1427-REIS_1423) as the query. *Bottom*, a portion of the large protein (PFX12133, 2285 aa) from the coral *Stylophora pistillata* shares similarity with these smaller *Rickettsia* proteins. (**B**) rCRCT-2 proteins are analogous to rCRCT-1 proteins. Alignment performed using MUSCLE with default settings (Edgar, 2004). (**C**) rCRCA-3 proteins recovered from Blastp searches using *R*. *bellii* rCRCA-3b as a query (the largest rCRCA-3 protein). na, sequences not recovered in Blastp searches against the NCBI ‘Rickettsia’ database but retrieved from PATRIC (asterisks denote PATRIC Local Family IDs). Subjects are listed accordingly to their placement in the phylogeny presented in **Figure 5E**. Colored boxes unite rCRCA-3 proteins recombined within the same genome. (**D**) rCRCT-3a HaloBalst analysis. *R*. *tamurae* rCRCT-3a (WP_051965318) was used as the query. Concentric halos depict hierarchical taxonomic databases increasing in divergence from the center. Average *Sm* score (see text for details) for all subjects and top ten subjects are provided, with highest score per database highlighted. ‘na’, not applicable. (**E**) Top 25 subjects from the Blastp search against ‘Proteobacteria’ using *R*. *tamurae* rCRCT-3a as the query. (**F**) Comparison of the *Cupriavidus taiwanensis* str. DSM 17343 CdiA toxin (Uniprot acc. B3R1C1) to rCRCT-3a of *R*. *tamurae*. Domains were predicted with SMART (Letunic and Bork, 2017). Amino acid similarity (%ID, red shading) was assessed using Blastp. (**G**) Structural analysis of a rCRCT/CRCA-3a module. (***top***) Alignment of residues 3391-3469 of *C*. *taiwanensis* CdiA, *R*. *tamurae* rCRCT-3a, and REIP rCRCT-3a. Structural information from *C*. *taiwanensis* CdiA (PDB:5T87) (Kryshtafovych et al., 2018) is provided at top. Alignment performed using MUSCLE with default settings (Edgar, 2004). (***bottom left***) Modeling with Phyre2 of *R*. *tamurae* rCRCT-3a to the CdiA-CT toxin structure of *C*. *taiwanensis* CdiA (PDB:5T87). The threading was done with 100% confidence. (***bottom right***) While the *C*. *taiwanensis* module was solved as a co-complex (Kryshtafovych et al., 2018), Phyre2 modeling could not thread *R*. *tamurae* rCRCA-3a to the antidote CdiI within the TA co-complex. The best model (46.3% confidence, 12% ID) for *R*. *tamurae* rCRCA-3a was to the structure of *Drosophila melanogaster* MAST/Orbit N-terminal domain PDB:4G3A (De La Mora-Rey et al., 2013)

**FIGURE S5.**
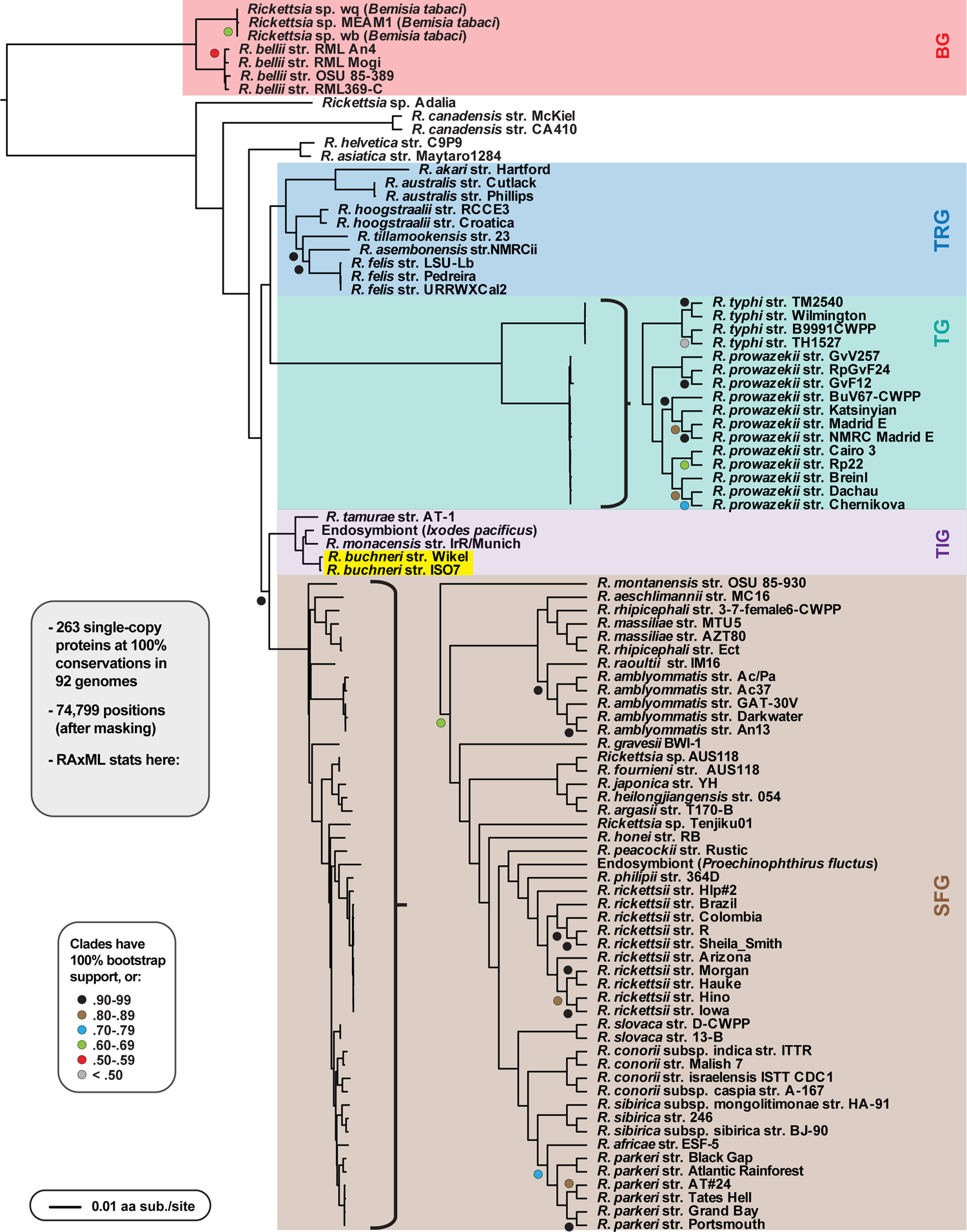
*Rickettsia* genome-based phylogeny estimation. *Rickettsia* groups follow previous classification (Gillespie et al., 2007), except that we recognize Tamurae/Ixodes Group (TIG) rickettsiae as a distinct clade from SFG rickettsiae. *R. buchneri* is highlighted. Phylogeny was estimated for 92 *Rickettsia* genomes; gray inset described details (see “Materials and Methods” for more details). Branch support was assessed with 1,000 pseudoreplications.

**FIGURE S6.**
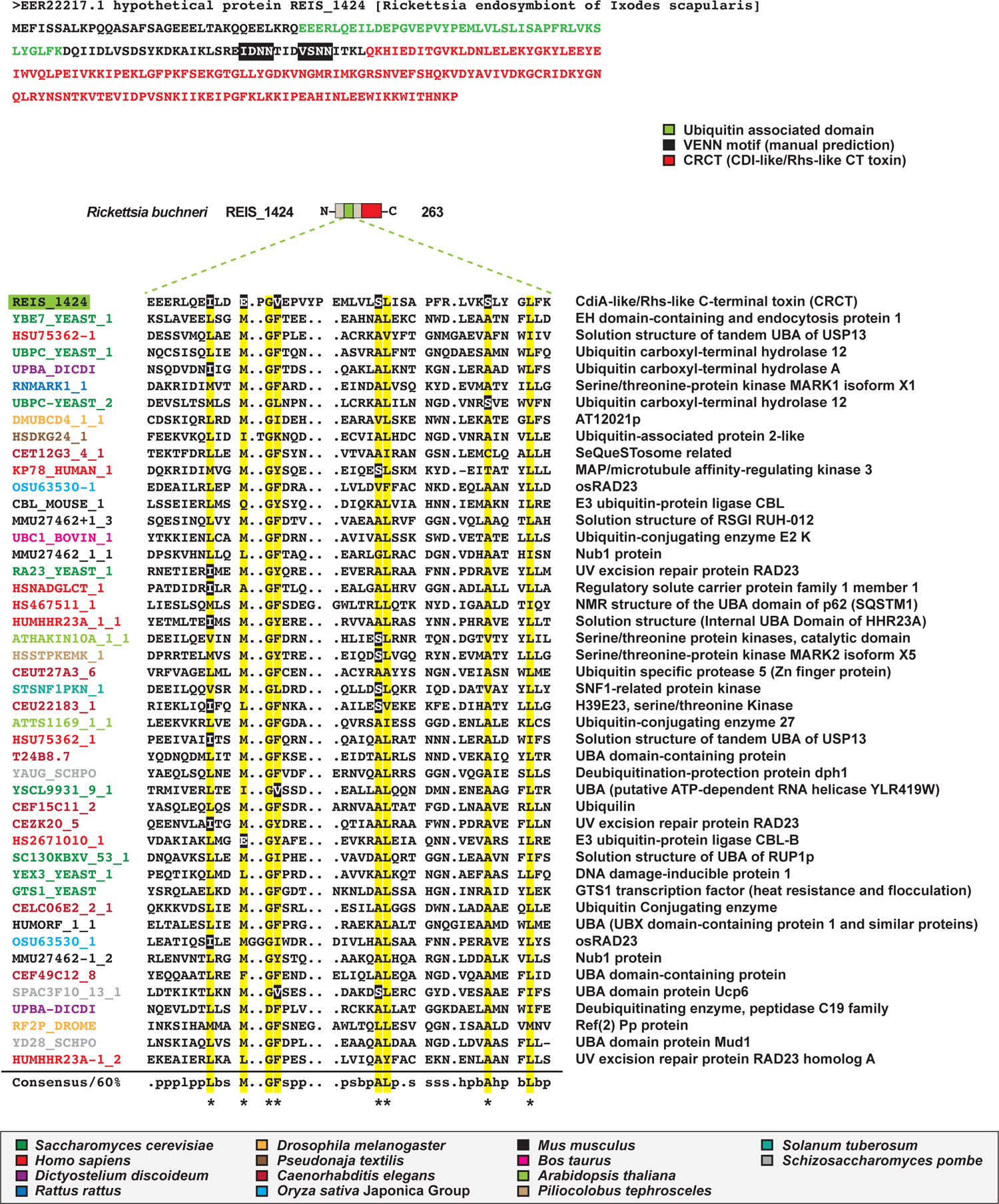
REIS_1424 contains a ubiquitin-associated domain demarcated from the C-terminal toxin by two predicted VENN motifs. (Mueller and Feigon, 2002); (E) Schemas for a few. Accession nos. Described domains here or in the M&M and get refs from SMART (Letunic and Bork, 2017). Wolbachia supergroups are within colored ellipses. Ctenocephalides felis-

**FIGURE S7.**
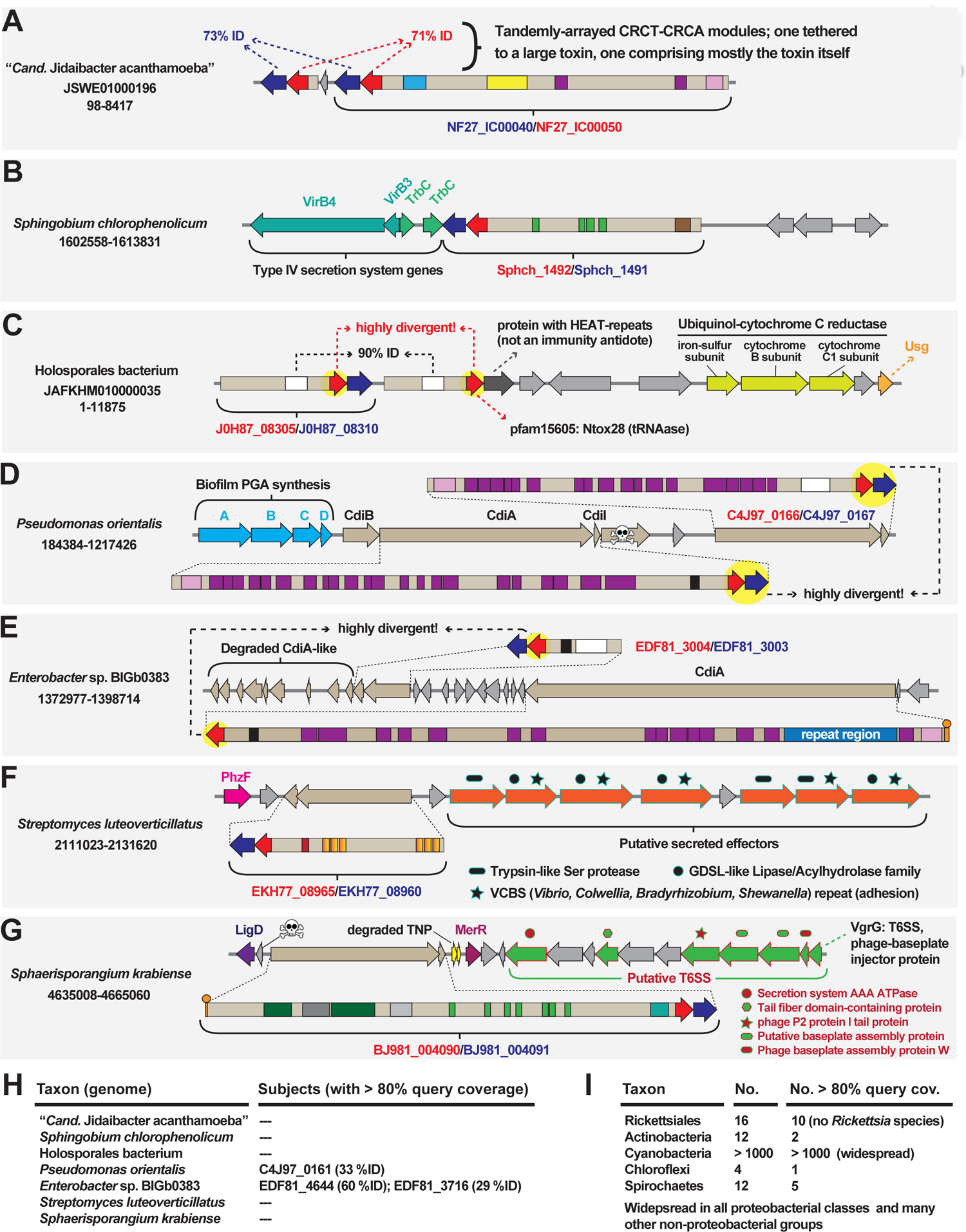
Gene neighborhoods for CRCT/CRCA modules in diverse bacteria. (**A-G**) Genome neighborhoods for the CRCT/CRCA modules shown in Figures 1 and 2 for (**A**) “*Candidatus* Jidaibacter acanthamoeba”, (**B**) *Sphingobium chlorophenolicum*, (**C**) Holosporales bacterium, (**D**) *Pseudomonas orientalis*, (**E**) *Enterobacter* sp. BIGb0383, *Streptomyces luteoverticillatus*, (**G**) *Sphaerisporangium krabiense*. (**H**) Results of Blastp searches against these seven genomes (specific taxon databases at NCBI) using the CdiB protein of *Escherichia coli* (UniProtKB-Q3YL97). (**I**) Occurrence of CdiB and CdiB-like proteins in the major taxa in which CRCT/CRCA modules were identified.

## References

1. Abraham, N. M., Liu, L., Jutras, B. L., Yadav, A. K., Narasimhan, S., Gopalakrishnan, V., et al. (2017). Pathogen-mediated manipulation of arthropod microbiota to promote infection. Proc. Natl. Acad. Sci. 114, E781–E790. doi:10.1073/pnas.1613422114.

2. Al-Khafaji, A. M., Armstrong, S. D., Varotto Boccazzi, I., Gaiarsa, S., Sinha, A., Li, Z., et al. (2020). Rickettsia buchneri, symbiont of the deer tick Ixodes scapularis, can colonise the salivary glands of its host. Ticks Tick. Borne. Dis. 11, 101299. doi:10.1016/J.TTBDIS.2019.101299.

3. Aoki, S. K., Diner, E. J., De Roodenbeke, C. T. K., Burgess, B. R., Poole, S. J., Braaten, B. A., et al. (2010). A widespread family of polymorphic contact-dependent toxin delivery systems in bacteria. Nat. 2010 *4687322* 468, 439–442. doi:10.1038/nature09490.

4. Aoki, S. K., Webb, J. S., Braaten, B. A., and Low, D. A. (2009). Contact-dependent growth inhibition causes reversible metabolic downregulation in Escherichia coli. J. Bacteriol. 191, 1777–1786. doi:10.1128/JB.01437-08/ASSET/94ABFA58-D59C-4950-AC3A-44EED9C05FEE/ASSETS/GRAPHIC/ZJB0060985500007.JPEG.

5. Aziz, R. K., Bartels, D., Best, A. A., DeJongh, M., Disz, T., Edwards, R. A., et al. (2008). The RAST Server: Rapid Annotations using Subsystems Technology. BMC Genomics 9, 75. doi:10.1186/1471-2164-9-75.

6. Balvín, O., Roth, S., Talbot, B., and Reinhardt, K. (2018). Co-speciation in bedbug Wolbachia parallel the pattern in nematode hosts. Sci. Rep. 8, 8797. doi:10.1038/s41598-018-25545-y.

7. Beck, C. M., Morse, R. P., Cunningham, D. A., Iniguez, A., Low, D. A., Goulding, C. W., et al. (2014). CdiA from enterobacter cloacae delivers a toxic ribosomal RNase into target bacteria. Structure 22, 707–718. doi:10.1016/J.STR.2014.02.012/ATTACHMENT/638214C1-C3F9-493D-BF50-00C4042D7460/MMC1.PDF.

8. Beckmann, J. F., Ronau, J. A., and Hochstrasser, M. (2017). A Wolbachia deubiquitylating enzyme induces cytoplasmic incompatibility. Nat. Microbiol. 2, 17007. doi:10.1038/nmicrobiol.2017.7.

9. Benson, M. J., Gawronski, J. D., Eveleigh, D. E., and Benson, D. R. (2004). Intracellular Symbionts and Other Bacteria Associated with Deer Ticks (Ixodes scapularis) from Nantucket and Wellfleet, Cape Cod, Massachusetts. Appl. Environ. Microbiol. 70, 616–620. doi:10.1128/AEM.70.1.616-620.2004.

10. Billings, A. N., Teltow, G. J., Weaver, S. C., and Walker, D. H. Molecular characterization of a novel Rickettsia species from Ixodes scapularis in Texas. Emerg. Infect. Dis. 4, 305–9. doi:10.3201/eid0402.980221.

11. Blanc, G., Ogata, H., Robert, C., Audic, S., Claverie, J. M., and Raoult, D. (2007). Lateral gene transfer between obligate intracellular bacteria: Evidence from the Rickettsia massiliae genome. Genome Res. 17, 1657–1664. doi:10.1101/gr.6742107.

12. Bruner, S. D., Norman, D. P. G., and Verdine, G. L. (2000). Structural basis for recognition and repair of the endogenous mutagen 8-oxoguanine in DNA. Nat. 2000 *4036772* 403, 859–866. doi:10.1038/35002510.

13. Burgdorfer, W., Hayes, S. F., and Mavros, A. J. (1981). “Rickettsiae and rickettsial diseases,” in, eds. W. Burgdorfer and R. L. Anacker (New York, NY.: Academic Press), 585–594. doi:10.3/JQUERY-UI.JS.

14. Busby, J. N., Panjikar, S., Landsberg, M. J., Hurst, M. R. H., and Lott, J. S. (2013). The BC component of ABC toxins is an RHS-repeat-containing protein encapsulation device. Nature 501, 547–50. doi:10.1038/nature12465.

15. Crooks, G. E., Hon, G., Chandonia, J.-M., and Brenner, S. E. (2004). WebLogo: a sequence logo generator. Genome Res. 14, 1188–90. doi:10.1101/gr.849004.

16. Cross, S. T., Kapuscinski, M. L., Perino, J., Maertens, B. L., Weger-Lucarelli, J., Ebel, G. D., et al. (2018). Co-Infection Patterns in Individual Ixodes scapularis Ticks Reveal Associations between Viral, Eukaryotic and Bacterial Microorganisms. Viruses 10. doi:10.3390/v10070388.

17. Cull, B., Burkhardt, N. Y., Wang, X. R., Thorpe, C. J., Oliver, J. D., Kurtti, T. J., et al. (2022). The Ixodes scapularis Symbiont Rickettsia buchneri Inhibits Growth of Pathogenic Rickettsiaceae in Tick Cells: Implications for Vector Competence. Front. Vet. Sci. 8, 1516. doi:10.3389/FVETS.2021.748427/BIBTEX.

18. Darby, A. C., Cho, N.-H., Fuxelius, H.-H., Westberg, J., and Andersson, S. G. E. (2007). Intracellular pathogens go extreme: genome evolution in the Rickettsiales. Trends Genet. 23, 511–520. doi:10.1016/j.tig.2007.08.002.

19. De La Mora-Rey, T., Guenther, B. D., and Finzel, B. C. (2013). The structure of the TOG-like domain of Drosophila melanogaster Mast/Orbit. urn:issn: 1744-3091 69, 723–729. doi:10.1107/S1744309113015182.

20. Driscoll, T. P., Gillespie, J. J., Nordberg, E. K., Azad, A. F., and Sobral, B. W. (2013). Bacterial DNA sifted from the Trichoplax adhaerens (Animalia: Placozoa) genome project reveals a putative rickettsial endosymbiont. Genome Biol. Evol. 5, 621–45. doi:10.1093/gbe/evt036.

21. Driscoll, T. P., Verhoeve, V. I., Brockway, C., Shrewsberry, D. L., Plumer, M., Sevdalis, S. E., et al. (2020). Evolution of *Wolbachia* mutualism and reproductive parasitism: insight from two novel strains that co-infect cat fleas. PeerJ 8, e10646. doi:10.7717/peerj.10646.

22. Edgar, R. C. (2004). MUSCLE: Multiple sequence alignment with high accuracy and high throughput. Nucleic Acids Res. 32, 1792–1797. doi:10.1093/nar/gkh340.

23. Feris, K. P., Ramsey, P. W., Frazar, C., Rillig, M. C., and Gannon, J. E. (2003). Structure and seasonal dynamics of hyporheic zone microbial communities in free-stone rivers of the estern United States. Microb. Ecol.

24. Finn, R. D., Clements, J., and Eddy, S. R. (2011). HMMER web server: interactive sequence similarity searching. Nucleic Acids Res. 39, W29–W37. doi:10.1093/NAR/GKR367.

25. Fuxelius, H.-H., Darby, A. C., Cho, N.-H., and Andersson, S. G. (2008). Visualization of pseudogenes in intracellular bacteria reveals the different tracks to gene destruction. Genome Biol. 2008 *92* 9, 1–15. doi:10.1186/GB-2008-9-2-R42.

26. Gerth, M., and Bleidorn, C. (2017). Comparative genomics provides a timeframe for Wolbachia evolution and exposes a recent biotin synthesis operon transfer. Nat. Microbiol. 2, 16241. doi:10.1038/nmicrobiol.2016.241.

27. Gil, J. C., Helal, Z. H., Risatti, G., and Hird, S. M. (2020). Ixodes scapularis microbiome correlates with life stage, not the presence of human pathogens, in ticks submitted for diagnostic testing. PeerJ 8, e10424. doi:10.7717/PEERJ.10424/SUPP-10.

28. Gillespie, J. J., Beier, M. S., Rahman, M. S., Ammerman, N. C., Shallom, J. M., Purkayastha, A., et al. (2007). Plasmids and rickettsial evolution: insight from Rickettsia felis. PLoS One 2, e266. doi:10.1371/journal.pone.0000266.

29. Gillespie, J. J., Driscoll, T. P., Verhoeve, V. I., Rahman, M. S., Macaluso, K. R., and Azad, A. F. (2018). A Tangled Web: Origins of Reproductive Parasitism. Genome Biol. Evol. 10, 2292–2309. doi:10.1093/gbe/evy159.

30. Gillespie, J. J., Driscoll, T. P., Verhoeve, V. I., Utsuki, T., Husseneder, C., Chouljenko, V. N., et al. (2015a). Genomic Diversification in Strains of Rickettsia felis Isolated from Different Arthropods. Genome Biol. Evol. 7, 35–56. doi:10.1093/gbe/evu262.

31. Gillespie, J. J., Joardar, V., Williams, K. P., Driscoll, T. P., Hostetler, J. B., Nordberg, E., et al. (2012a). A Rickettsia genome overrun by mobile genetic elements provides insight into the acquisition of genes characteristic of an obligate intracellular lifestyle. J. Bacteriol. 194, 376–94. doi:10.1128/JB.06244-11.

32. Gillespie, J. J., Kaur, S. J., Sayeedur Rahman, M., Rennoll-Bankert, K., Sears, K. T., Beier-Sexton, M., et al. (2015b). Secretome of obligate intracellular Rickettsia. FEMS Microbiol. Rev. 39. doi:10.1111/1574-6976.12084.

33. Gillespie, J. J., Nordberg, E. K., Azad, A. A., and Sobral, B. W. (2012b). “Phylogeny And Comparative Genomics: The Shifting Landscape In The Genomics Era,” in Intracellular Pathogens II: Rickettsiales, ed. A.F. Azad and G.H. Palmer (Boston: American Society of Microbiology), 84–141.

34. Gillespie, J. J., Williams, K., Shukla, M., Snyder, E. E., Nordberg, E. K., Ceraul, S. M., et al. (2008). Rickettsia phylogenomics: Unwinding the intricacies of obligate intracellular life. PLoS One 3. doi:10.1371/journal.pone.0002018.

35. Gulia-Nuss, M., Nuss, A. B. A. B., Meyer, J. M. J. M., Sonenshine, D. E. D. E., Roe, R. M. M., Waterhouse, R. M. R. M., et al. (2016). Genomic insights into the Ixodes scapularis tick vector of Lyme disease. Nat. Commun. 7, 10507. doi:10.1038/ncomms10507.

36. Hagen, R., Verhoeve, V. I., Gillespie, J. J., and Driscoll, T. P. (2018). Conjugative transposons and their cargo genes vary across natural populations of Rickettsia buchneri infecting the tick Ixodes scapularis. Genome Biol. Evol. 10, 3218–3229. doi:10.1093/gbe/evy247.

37. Johnson, P. M., Gucinski, G. C., Garza-Sánchez, F., Wong, T., Hung, L. W., Hayes, C. S., et al. (2016). Functional Diversity of Cytotoxic tRNase/Immunity Protein Complexes from Burkholderia pseudomallei*. J. Biol. Chem. 291, 19387–19400. doi:10.1074/JBC.M116.736074.

38. Ju, J.-F., Bing, X.-L., Zhao, D.-S., Guo, Y., Xi, Z., Hoffmann, A. A., et al. (2019). Wolbachia supplement biotin and riboflavin to enhance reproduction in planthoppers. ISME J., 1–12. doi:10.1038/s41396-019-0559-9.

39. Kajava, A. V., Cheng, N., Cleaver, R., Kessel, M., Simon, M. N., Willery, E., et al. (2001). Beta-helix model for the filamentous haemagglutinin adhesin of Bordetella pertussis and related bacterial secretory proteins. Mol. Microbiol. 42, 279–292. doi:10.1046/J.1365-2958.2001.02598.X.

40. Kelley, L. A., and Sternberg, M. J. E. (2009). Protein structure prediction on the Web: a case study using the Phyre server. Nat. Protoc. 4, 363–371. doi:10.1038/nprot.2009.2.

41. Kryshtafovych, A., Albrecht, R., Baslé, A., Bule, P., Caputo, A. T., Carvalho, A. L., et al. (2018). Target highlights from the first post-PSI CASP experiment (CASP12, May– August 2016). Proteins Struct. Funct. Bioinforma. 86, 27–50. doi:10.1002/PROT.25392.

42. Kurtti, T. J., Burkhardt, N. Y., Heu, C. C., and Munderloh, U. G. (2016). Fluorescent Protein Expressing Rickettsia buchneri and Rickettsia peacockii for Tracking Symbiont-Tick Cell Interactions. Vet. Sci. 2016, Vol. 3, Page 34 3, 34. doi:10.3390/VETSCI3040034.

43. Kurtti, T. J., Felsheim, R. F., Burkhardt, N. Y., Oliver, J. D., Heu, C. C., and Munderloh, U. G. (2015). Rickettsia buchneri sp. nov., a rickettsial endosymbiont of the blacklegged tick Ixodes scapularis. Int. J. Syst. Evol. Microbiol. 65, 965–70. doi:10.1099/ijs.0.000047.

44. Lee, S., Kakumanu, M. L., Ponnusamy, L., Vaughn, M., Funkhouser, S., Thornton, H., et al. (2014). The skin of outdoor workers in North Carolina. Parasites and Vectors 7. doi:10.1186/s13071-014-0607-2.

45. Lehane, M. J., and Lehane, M. J. (2010). “Managing the blood meal,” in The Biology of Blood-Sucking in Insects (Cambridge University Press), 84–115. doi:10.1017/cbo9780511610493.007.

46. LePage, D. P., Metcalf, J. A., Bordenstein, S. R., On, J., Perlmutter, J. I., Shropshire, J. D., et al. (2017). Prophage WO genes recapitulate and enhance Wolbachia-induced cytoplasmic incompatibility. Nature 543, 243–247. doi:10.1038/nature21391.

47. Letunic, I., and Bork, P. (2017). 20 years of the SMART protein domain annotation resource. Nucleic Acids Res. doi:10.1093/nar/gkx922.

48. Levin, M. L., Schumacher, L. B. M., and Snellgrove, A. (2018). Effects of Rickettsia amblyommatis Infection on the Vector Competence of Amblyomma americanum Ticks for Rickettsia rickettsii. https://home.liebertpub.com/vbz 18, 579–587. doi:10.1089/VBZ.2018.2284.

49. Lin, H. H., Filloux, A., and Lai, E. M. (2020). Role of Recipient Susceptibility Factors During Contact-Dependent Interbacterial Competition. Front. Microbiol. 11, 2768. doi:10.3389/FMICB.2020.603652/BIBTEX.

50. Liu, W., Xie, Y., Ma, J., Luo, X., Nie, P., Zuo, Z., et al. (2015). IBS: an illustrator for the presentation and visualization of biological sequences. Bioinformatics 31, 3359–61. doi:10.1093/bioinformatics/btv362.

51. Liu, X., Bulgakov, O. V., Darrow, K. N., Pawlyk, B., Adamian, M., Liberman, M. C., et al. (2007). Usherin is required for maintenance of retinal photoreceptors and normal development of cochlear hair cells. Proc. Natl. Acad. Sci. 104, 4413–4418. doi:10.1073/PNAS.0610950104.

52. Macaluso, K. R., Sonenshine, D. E., Ceraul, S. M., and Azad, A. F. (2002). Rickettsial Infection in Dermacentor variabilis (Acari: Ixodidae) Inhibits Transovarial Transmission of a Second Rickettsia. J. Med. Entomol. 39, 809–813. doi:10.1603/0022-2585-39.6.809.

53. Madison-Antenucci, S., Kramer, L. D., Gebhardt, L. L., and Kauffman, E. (2020). Emerging Tick-Borne Diseases. Clin. Microbiol. Rev. 33. doi:10.1128/CMR.00083-18.

54. Magnarelli, L. A., Andreadis, T. G., Stafford, K. C., and Holland, C. J. (1991). Rickettsiae and Borrelia burgdorferi in ixodid ticks. J. Clin. Microbiol. 29, 2798– 2804.

55. Manzano-Marín, A., Oceguera-Figueroa, A., Latorre, A., Jiménez-García, L. F., and Moya, A. (2015). Solving a bloody mess: B-vitamin independentmetabolic convergence among gammaproteobacterial obligate endosymbionts from blood-feeding arthropods and the Leech haementeria officinalis. Genome Biol. Evol. 7, 2871–2884. doi:10.1093/gbe/evv188.

56. Maurin, M., and Raoult, D. (2001). Use of Aminoglycosides in Treatment of Infections Due to Intracellular Bacteria. Antimicrob. Agents Chemother. 45, 2977–2986. doi:10.1128/AAC.45.11.2977-2986.2001.

57. Michalska, K., Quan Nhan, D., Willett, J. L. E., Stols, L. M., Eschenfeldt, W. H., Jones, A. M., et al. (2018). Functional plasticity of antibacterial EndoU toxins. Mol. Microbiol. 109, 509–527. doi:10.1111/MMI.14007.

58. Miller, D. L., Smith, E. A., and Newton, I. L. G. (2020). A bacterial symbiont protects honey bees from fungal disease. bioRxiv, 2020.01.21.914325. doi:10.1101/2020.01.21.914325.

59. Moreno, C. X., Moy, F., Daniels, T. J., Godfrey, H. P., and Cabello, F. C. (2006). Molecular analysis of microbial communities identified in different developmental stages of Ixodes scapularis ticks from Westchester and Dutchess Counties, New York. Environ. Microbiol. 8, 761–72. doi:10.1111/j.1462-2920.2005.00955.x.

60. Morse, R. P., Nikolakakis, K. C., Willett, J. L. E., Gerrick, E., Low, D. A., Hayes, C. S., et al. (2012). Structural basis of toxicity and immunity in contact-dependent growth inhibition (CDI) systems. Proc. Natl. Acad. Sci. U. S. A. 109, 21480–21485. doi:10.1073/PNAS.1216238110/-/DCSUPPLEMENTAL.

61. Mueller, T. D., and Feigon, J. (2002). Solution Structures of UBA Domains Reveal a Conserved Hydrophobic Surface for Protein–Protein Interactions. J. Mol. Biol. 319, 1243–1255. doi:10.1016/S0022-2836(02)00302-9.

62. Munderloh, U. G., Jauron, S. D., Fingerle, V., Leitritz, L., Hayes, S. F., Hautman, J. M., et al. (1999). Invasion and intracellular development of the human granulocytic ehrlichiosis agent in tick cell culture. J. Clin. Microbiol. 37, 2518–2524. doi:10.1128/JCM.37.8.2518-2524.1999/ASSET/225E7731-33D8-4A77-AB6E-0910DAE543E2/ASSETS/GRAPHIC/JM0890090005.JPEG.

63. Munderloh, U. G., Jauron, S. D., and Kurtti, T. J. (2005). “Chapter 3: The Tick: a Different Kind of Host for Human Pathogens,” in Tick-Borne Diseases of Humans, eds. J. L. Goodman, D. T. Dennis, and D. E. Sonenshine (Boston: American Society for Microbiology).

64. Narasimhan, S., Rajeevan, N., Liu, L., Zhao, Y. O., Heisig, J., Pan, J., et al. (2014). Gut Microbiota of the Tick Vector Ixodes scapularis Modulate Colonization of the Lyme Disease Spirochete. Cell Host Microbe 15, 58–71. doi:10.1016/j.chom.2013.12.001.

65. Nikoh, N., Hosokawa, T., Moriyama, M., Oshima, K., Hattori, M., and Fukatsu, T. (2014). Evolutionary origin of insect-Wolbachia nutritional mutualism. Proc. Natl. Acad. Sci. U. S. A. 111, 10257–62. doi:10.1073/pnas.1409284111.

66. Nikolakakis, K., Amber, S., Wilbur, J. S., Diner, E. J., Aoki, S. K., Poole, S. J., et al. (2012). The toxin/immunity network of Burkholderia pseudomallei contact-dependent growth inhibition (CDI) systems. Mol. Microbiol. 84, 516. doi:10.1111/J.1365-2958.2012.08039.X.

67. Ogata, H., La Scola, B., Audic, S., Renesto, P., Blanc, G., Robert, C., et al. (2006). Genome sequence of Rickettsia bellii illuminates the role of amoebae in gene exchanges between intracellular pathogens. PLoS Genet. 2, 733–744. doi:10.1371/JOURNAL.PGEN.0020076.

68. Oliver, J. D., Price, L. D., Burkhardt, N. Y., Heu, C. C., Khoo, B. S., Thorpe, C. J., et al. (2021). Growth Dynamics and Antibiotic Elimination of Symbiotic Rickettsia buchneri in the Tick Ixodes scapularis (Acari: Ixodidae). Appl. Environ. Microbiol. 87, 1–9. doi:10.1128/AEM.01672-20.

69. Penz, T., Schmitz-Esser, S., Kelly, S. E., Cass, B. N., Müller, A., Woyke, T., et al. (2012). Comparative Genomics Suggests an Independent Origin of Cytoplasmic Incompatibility in Cardinium hertigii. PLoS Genet. 8, e1003012. doi:10.1371/journal.pgen.1003012.

70. Perler, F. B. (1998). Protein Splicing of Inteins and Hedgehog Autoproteolysis: Structure, Function, and Evolution. Cell 92, 1–4. doi:10.1016/S0092-8674(00)80892-2.

71. Poole, S., Diner, E., Aoki, S., Braaten, B., t’Kint de Roodenbeke, C., Low, D., et al. (2011). Identification of functional toxin/immunity genes linked to contact-dependent growth inhibition (CDI) and rearrangement hotspot (Rhs) systems. PLoS Genet. 7. doi:10.1371/JOURNAL.PGEN.1002217.

72. Rawlings, N. D., and Barrett, A. J. (1995). Evolutionary families of metallopeptidases. Methods Enzymol. 248, 183–228. doi:10.1016/0076-6879(95)48015-3.

73. Ríhová, J., Nováková, E., Husník, F., and Hypša, V. (2017). Legionella Becoming a Mutualist: Adaptive Processes Shaping the Genome of Symbiont in the Louse Polyplax serrata. Genome Biol. Evol. 9, 2946–2957. doi:10.1093/gbe/evx217.

74. Rolain, J. M., Maurin, M., Vestris, G., and Raoult, D. (1998). In vitro susceptibilities of 27 rickettsiae to 13 antimicrobials. Antimicrob. Agents Chemother. 42, 1537–1541. doi:10.1128/AAC.42.7.1537.

75. Ross, B. D., Hayes, B., Radey, M. C., Lee, X., Josek, T., Bjork, J., et al. (2018). Ixodes scapularis does not harbor a stable midgut microbiome. ISME J. 12, 2596–2607. doi:10.1038/s41396-018-0161-6.

76. Ruhe, Z. C., Subramanian, P., Song, K., Nguyen, J. Y., Stevens, T. A., Low, D. A., et al. (2018). Programmed Secretion Arrest and Receptor-Triggered Toxin Export during Antibacterial Contact-Dependent Growth Inhibition. Cell 175, 921–933.e14. doi:10.1016/J.CELL.2018.10.033/ATTACHMENT/98B0A2F7-BB29-42E7-8C39-7312137CD8F5/MMC1.PDF.

77. Sanchez-Vicente, S., Tagliafierro, T., Coleman, J. L., Benach, J. L., and Tokarz, R. (2019). Polymicrobial nature of tick-borne diseases. MBio 10. doi:10.1128/mBio.02055-19.

78. Schulz, F., Martijn, J., Wascher, F., Lagkouvardos, I., Kostanjšek, R., Ettema, T. J. G., et al. (2016). A Rickettsiales symbiont of amoebae with ancient features. Environ. Microbiol. 18, 2326–2342. doi:10.1111/1462-2920.12881.

79. Simser, J. A., Palmer, A. T., Fingerle, V., Wilske, B., Kurtti, T. J., and Munderloh, U. G. (2002). Rickettsia monacensis sp. nov., a spotted fever group Rickettsia, from ticks (Ixodes ricinus) collected in a European city park. Appl. Environ. Microbiol. 68, 4559–66. Available at: http://www.ncbi.nlm.nih.gov/pubmed/12200314 [Accessed January 26, 2015].

80. Stamatakis, a, Ludwig, T., and Meier, H. (2005). RAxML-III: a fast program for maximum likelihood-based inference of large phylogenetic trees. Bioinformatics 21, 456–63. doi:10.1093/bioinformatics/bti191.

81. Stamatakis, A. (2014). RAxML version 8: A tool for phylogenetic analysis and post-analysis of large phylogenies. Bioinformatics 30, 1312–1313. doi:10.1093/bioinformatics/btu033.

82. Steiner, F. E., Pinger, R. R., Vann, C. N., Grindle, N., Civitello, D., Clay, K., et al. (2008). Infection and Co-infection Rates of Anaplasma phagocytophilum

83. Variants, Babesia spp., Borrelia burgdorferi, and the Rickettsial Endosymbiont in Ixodes scapularis (Acari: Ixodidae) from Sites in Indiana, Maine, Pennsylvania, and. J. Med. Entomol. 45, 289–297. doi:10.1603/0022-2585(2008)45[289:iacroa]2.0.co;2.

84. Stenos, J., Graves, S. R., and Unsworth, N. B. (2005). A highly sensitive and specific real-time PCR assay for the detection of spotted fever and typhus group rickettsiae. Am. J. Trop. Med. Hyg. 73, 1083–1085. doi:10.4269/ajtmh.2005.73.1083.

85. Swanson, K. I., and Norris, D. E. (2007). Co-circulating microorganisms in questing Ixodes scapularis nymphs in Maryland. J. Vector Ecol. 32, 243. doi:10.3376/1081-1710(2007)32[243:cmiqis]2.0.co;2.

86. Talavera, G., and Castresana, J. (2007). Improvement of phylogenies after removing divergent and ambiguously aligned blocks from protein sequence alignments. Syst. Biol. 56, 564–77. doi:10.1080/10635150701472164.

87. Thapa, S., Zhang, Y., and Allen, M. S. (2019). Bacterial microbiomes of Ixodes scapularis ticks collected from Massachusetts and Texas, USA. BMC Microbiol. 19. doi:10.1186/s12866-019-1514-7.

88. Tokarz, R., Tagliafierro, T., Sameroff, S., Cucura, D. M., Oleynik, A., Che, X., et al. (2019). Microbiome analysis of Ixodes scapularis ticks from New York and Connecticut. Ticks Tick. Borne. Dis. 10, 894–900. doi:10.1016/j.ttbdis.2019.04.011.

89. Troughton, D. R., and Levin, M. L. (2007). Life Cycles of Seven Ixodid Tick Species (Acari: Ixodidae) Under Standardized Laboratory Conditions. J. Med. Entomol. 44, 732–740. doi:10.1093/jmedent/44.5.732.

90. van Treuren, W., Ponnusamy, L., Brinkerhoff, R. J., Gonzalez, A., Parobek, C. M., Juliano, J. J., et al. (2015). Variation in the microbiota of Ixodes ticks with regard to geography, species, and sex. Appl. Environ. Microbiol. 81, 6200–6209. doi:10.1128/AEM.01562-15.

91. Walker, D. H., and Ismail, N. (2008). Emerging and re-emerging rickettsioses: endothelial cell infection and early disease events. Nat. Rev. Microbiol. 6, 375–86. doi:10.1038/nrmicro1866.

92. Weyer, K., Boldt, H. B., Poulsen, C. B., Kjaer-Sorensen, K., Gyrup, C., and Oxvig, C. (2007). A Substrate Specificity-determining Unit of Three Lin12-Notch Repeat Modules Is Formed in Trans within the Pappalysin-1 Dimer and Requires a Sequence Stretch C-terminal to the Third Module *. J. Biol. Chem. 282, 10988– 10999. doi:10.1074/JBC.M607903200.

93. Willett, J. L. E., Ruhe, Z. C., Goulding, C. W., Low, D. A., and Hayes, C. S. (2015). Contact-Dependent Growth Inhibition (CDI) and CdiB/CdiA Two-Partner Secretion Proteins. J. Mol. Biol. 427, 3754–3765. doi:10.1016/J.JMB.2015.09.010.

94. Wormser, G. P., Dattwyler, R. J., Shapiro, E. D., Halperin, J. J., Steere, A. C., Klempner, M. S., et al. (2006). The Clinical Assessment, Treatment, and Prevention of Lyme Disease, Human Granulocytic Anaplasmosis, and Babesiosis: Clinical Practice Guidelines by the Infectious Diseases Society of America. Clin. Infect. Dis. 43, 1089–1134. doi:10.1086/508667.

95. Wright, C. L., Sonenshine, D. E., Gaff, H. D., and Hynes, W. L. (2015). Rickettsia parkeri Transmission to Amblyomma americanum by Cofeeding with Amblyomma maculatum (Acari: Ixodidae) and Potential for Spillover. J. Med. Entomol. 52, 1090– 1095. doi:10.1093/JME/TJV086.

96. Yang, Q., Kučerová, Z., Perlman, S. J., Opit, G. P., Mockford, E. L., Behar, A., et al. (2015). Morphological and molecular characterization of a sexually reproducing colony of the booklouse Liposcelis bostrychophila (Psocodea: Liposcelididae) found in Arizona. Sci. Rep. 5, 10429. doi:10.1038/srep10429.

97. Yeats, C., Bentley, S., and Bateman, A. (2003). New knowledge from old: In silico discovery of novel protein domains in Streptomyces coelicolor. BMC Microbiol. 3, 1–20. doi:10.1186/1471-2180-3-3/FIGURES/13.

98. Zeng, Z., Fu, Y., Guo, D., Wu, Y., Ajayi, O. E., and Wu, Q. (2018). Bacterial endosymbiont Cardinium cSfur genome sequence provides insights for understanding the symbiotic relationship in Sogatella furcifera host. BMC Genomics 19, 688. doi:10.1186/s12864-018-5078-y.

99. Zhang, D., Iyer, L. M., and Aravind, L. (2011). A novel immunity system for bacterial nucleic acid degrading toxins and its recruitment in various eukaryotic and DNA viral systems. Nucleic Acids Res. 39, 4532–4552. doi:10.1093/NAR/GKR036.

100. Zolnik, C. P., Prill, R. J., Falco, R. C., Daniels, T. J., and Kolokotronis, S. O. (2016). Microbiome changes through ontogeny of a tick pathogen vector. Mol. Ecol. 25, 4963–4977. doi:10.1111/MEC.13832.

